# ZNF217 promotes receptor tyrosine kinase plasticity and AXL-ERK dependency in ovarian cancer

**DOI:** 10.64898/2026.09.11.750913

**Authors:** Megha J Pandya, Jessica Hoffman, Achuth Padmanabhan

## Abstract

ZNF217 is a potent oncogene that drives ovarian cancer progression and therapeutic resistance. We show that ZNF217 overexpression markedly increases ERBB2 levels in ovarian cancer cells, suggesting its potential as a biomarker to identify ovarian tumors that will respond to ERBB2-targeted therapeutics. Unexpectedly, ZNF217-high ovarian cancer cells exhibit resistance to multiple ERBB2 inhibitors, revealing a disconnect between receptor abundance and drug sensitivity. Mechanistically, ZNF217 drives chemoresistance by elevating the expression of several key receptor tyrosine kinases, most notably AXL, that activates MAPK signaling. Further, ZNF217 also upregulates ERK1/2 levels in ovarian cancer cells. This induces rewiring of downstream signaling, promoting an ERK-dominant state that sustains survival despite ERBB2 inhibition. Therapeutic targeting of the AXL–ERK axis reduced cell viability and metastatic potential in vitro, while suppressing tumor burden and prolonging survival in xenograft models. Thus, by elevating the expression of several key signaling receptors and their downstream effectors in ovarian cancer cells, ZNF217 establishes signaling plasticity that defines a novel mechanism of chemoresistance. These findings identify ZNF217 as a key determinant of signaling state and drug response and provides rationale to evaluate targeting AXL and ERK signaling in ZNF217-high ovarian tumors.

## INTRODUCTION

Metastatic ovarian cancer is a lethal disease with limited therapeutic options^1, 2^. Due to the ineffectiveness of current treatment strategies and emergence of chemoresistance, prognosis for ovarian cancer patients with metastatic disease remains extremely poor^1–3^. Overcoming chemoresistance in ovarian cancer is a critical unmet therapeutic challenge and is essential for improving clinical outcome. Achieving this goal will require identifying factors that drive chemoresistance and elucidating the mechanisms by which they enable cancer cells to evade and survive therapeutic interventions.

Recent reports establish the transcription factor Zinc Finger Protein 217 (ZNF217) as a potent oncogene in ovarian cancer cells^4^. In addition to promoting ovarian cancer metastasis, ZNF217 drives resistance to chemotherapeutic agents such as carboplatin, paclitaxel, and doxorubicin^4^. Consequently, ovarian cancer patients with elevated ZNF217 expression in their tumors have a poor prognosis^4^. Our efforts to identify therapeutic vulnerabilities in ZNF217 driven ovarian tumors identified significant upregulation in Human epidermal growth factor receptor 2 (ERBB2/HER2) in ZNF217-high ovarian cancer cells. ERBB2 is a receptor tyrosine kinase and a member of the epidermal growth factor receptor (EGFR) family^5^. Dysregulated ERBB2 expression is observed in several malignancies, such as breast and gastroesophageal cancers^6, 7^. Elevated ERBB2 functions as an oncogene in these tumors and promotes cancer progression and chemoresistance^7^. Although ERBB2 lacks a known ligand, it can heterodimerize with other ligand bound members of the EGFR family such as ERBB1 (EGFR/Her-1) and ERBB3/Her-3 to activate downstream signaling^8, 9^. In addition to members of the EGFR family, ERBB2 is also capable of interacting with other receptor tyrosine kinases such as members of the fibroblast growth factor receptor family, and AXL protein kinase to activate intracellular signaling pathways^10–12^. Ligand-independent activation of ERBB2 through homodimerization is also observed in cancer cells with elevated ERBB2 levels^13^. Notably, ERBB2 lacks a known ligand yet exhibits the most potent catalytic protein kinase activity among EGFR family members.

Activated ERBB2 can in fact signal either via the PI3K-AKT signaling pathway or the MAPK pathway in a context-dependent manner^14^. ERBB2 amplification and overexpression has been shown to activate PI3K-AKT signaling in breast cancer cells^15^. Although ERBB2 upregulation is associated with more aggressive disease in breast cancer patients, elevated ERBB2 levels serve as a biomarker to identify tumors that are likely to respond to ERBB2 targeting therapeutics^16^. Anti-ERBB2 drugs such as trastuzumab and lapatinib have been very effective in the treatment of ERBB2 overexpressing breast and gastroesophageal tumors^16^. Unlike breast and gastroesophageal cancers, ERBB2’s role or therapeutic utility in ovarian cancer is less understood. Studies show that ERBB2 expression at the protein level increases as ovarian cancer progresses, suggesting a potential role in metastasis^17^. As over 80 percent of ovarian cancer patients are diagnosed with advanced metastatic disease, ERBB2 can potentially impact prognosis in these patients^18^. Importantly, ERBB2 positive ovarian cancer cells are sensitive to trastuzumab (Herceptin), implying its utility as a therapeutic strategy in ERBB2-high metastatic ovarian tumors^17^. These data, together with the resounding success of anti-ERBB2 therapeutics, have generated significant interest in exploring ERBB2 as a therapeutic target in ovarian cancer^19,20^.

The effectiveness of ERBB2-targeting strategies will depend heavily on reliable biomarkers that will enable identification of potential responders. As ZNF217 upregulates ERBB2 in ovarian cancer cells, we hypothesized that ZNF217-driven tumors will be more sensitive to ERBB2 targeting drugs. However, despite elevated ERBB2 levels, ZNF217 overexpressing ovarian cancer cells were more resistant to anti-ERBB2 therapeutics. This paradoxical finding underscores the importance of understanding mechanisms that drives chemoresistance in ZNF217 overexpressing ovarian tumors. Our data reveal that ZNF217 causes upregulation of multiple receptor tyrosine kinases in addition to ERBB2, particularly AXL. This establishes signaling plasticity in ZNF217-high ovarian cancer cells, enabling them to rewire cellular signaling and resist chemotherapeutics more effectively. We show that ZNF217 overexpressing cells are dependent on both AXL and MAPK signaling pathway to drive oncogenic phenotypes. These results suggest that targeting these pathways will be an effective therapeutic strategy for ZNF217 driven ovarian tumors.

## RESULTS

### ZNF217 upregulates ERBB2 expression in ovarian cancer cells

The transcription factor ZNF217 is known to function as a potent oncogene in ovarian cancer cells^4^. As reported previously, stable ZNF217 overexpression drives an increase in cell viability and proliferation in ovarian cancer cells (Figs. 1A and 1B). RNAseq data from OVCA420 and TYK-Nu cell lines reveal that several processes that regulate cell proliferation and metastatic potential is altered upon ZNF217 overexpression (Fig. 1C). In addition to these processes, ERBB2 signaling is also altered upon ZNF217 overexpression in ovarian cancer cells (Fig. 1C). Similar results are observed using RNAseq data from OVCA420 cells in which ZNF217 is knocked down (Fig. S1A). Interestingly, RT-qPCR analysis shows that ZNF217 overexpression causes a 12-fold increase in ERBB2 mRNA levels in OVCA420 cells (Fig. 1D). Consistent with elevated ERBB2 mRNA levels, ectopic ZNF217 overexpression in OVCA420 and TYK-Nu cells resulted in a dramatic increase in ERBB2 protein levels (Fig. 1E). To further confirm the link between ZNF217 and ERBB2 in ovarian cancer cells, we determined the effect of ZNF217 depletion on ERBB2 levels. OVCA420 cells were engineered to stably express a doxycycline-inducible ZNF217 shRNA and doxycycline-inducible ZNF217 depletion in these cells was confirmed by western blot (Fig. IF). ZNF217 knockdown resulted in a reduction in ERBB2 level in these cells (Fig. 1G). Consistent with our data, analysis of publicly available gene expression data show that both ZNF217 and ERBB2 are elevated in ovarian tumors compared to normal tissue (Fig. 1H). These data show that ZNF217 is a positive regulator of ERBB2 levels in ovarian cancer cells.

**Figure 1.**
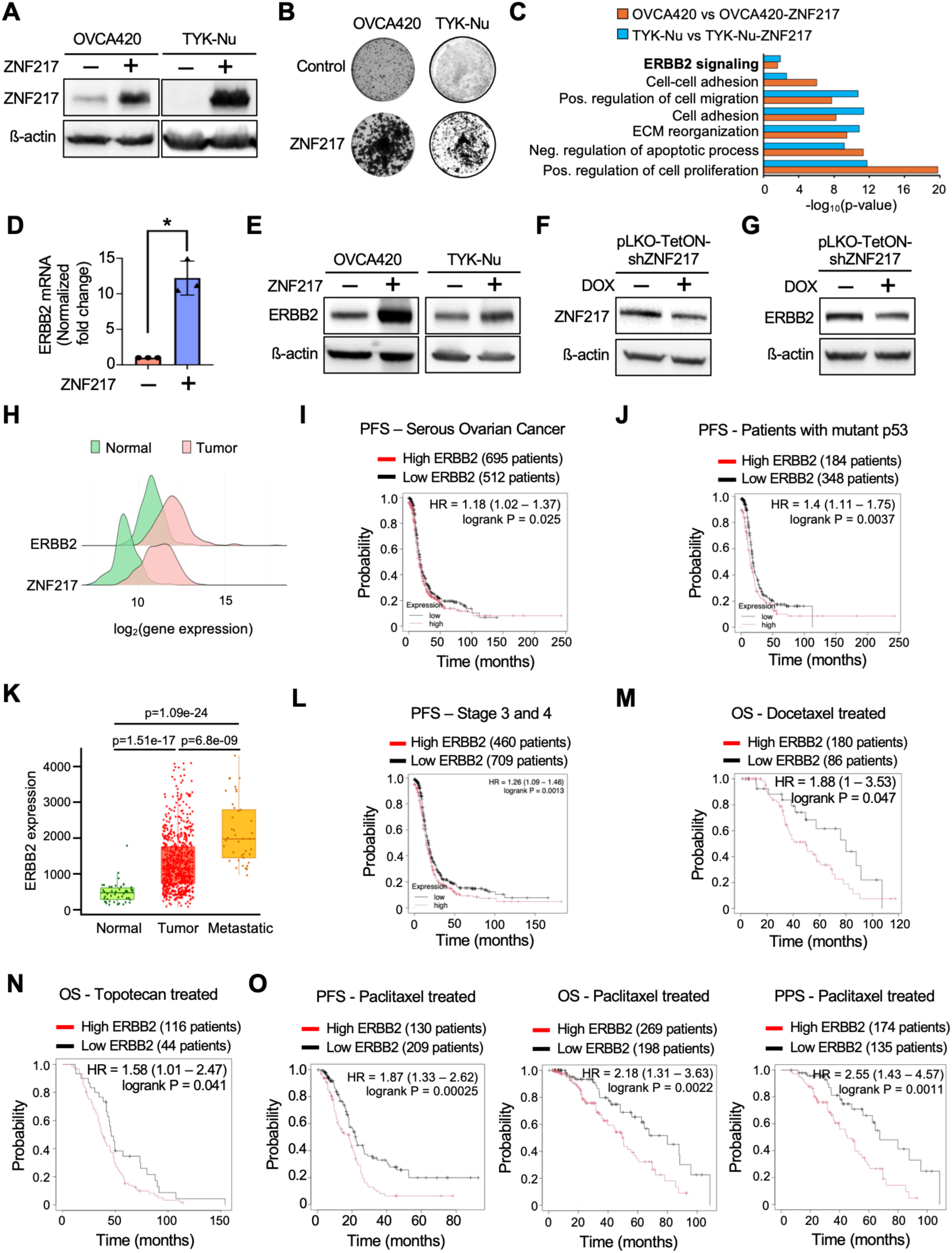
ZNF217 upregulates ERBB2 expression in ovarian cancer cells. **(A)** Western blots showing stable ZNF217 overexpression in OVCA420 and TYK-Nu cells. **(B)** Crystal violet staining reveals increased cell viability in ZNF217 overexpressing OVCA420 and TYK-Nu cells. **(C)**. Gene Ontology terms that are significantly enriched in ZNF217 overexpressing OVCA420 and TYK-Nu cells include ERBB2 signaling. **(D)** RT-qPCR data showing ERBB2 mRNA is upregulated in OVCA420-ZNF217 cells. **(E)** Western blots reveal ERBB2 upregulation in OVCA420-ZNF217 and TYK-Nu-ZNF217 cells. **(F)** ZNF217 levels are depleted upon Dox addition in OVCA420-Tet-ON-shZNF217 cells. **(G)** ZNF217 depletion decreases ERBB2 level in OVCA420 cells. **(H)** Expression of both ZNF217 and ERBB2 are elevated in ovarian tumors. Data was retrieved from TNMplot^21^. **(I)** Data from Kmplotter database for serous ovarian tumors show that patients with ERBB2-high tumors have lower progression-free survival (PFS)^22^. **(J)** Data from Kmplotter database for ovarian tumors with mutant p53 show that ERBB2-high tumors have lower progression-free survival (PFS)^22^. **(K)** ERBB2 expression is elevated in metastatic ovarian tumors. **(L)** ERBB2-high metastatic ovarian tumors have lower progression-free survival (PFS)^22^. **(M** and **N)** Ovarian cancer patients with ERBB2-high tumors exhibit lower overall survival (OS) when treated with **(M)** docetaxel and **(N)** topotecan. **(O)** Ovarian cancer patients with ERBB2-high tumors exhibit lower progression free survival (PFS), overall survival (OS), and post-progression survival (PPS) when treated with paclitaxel.

### ERBB2 expression correlates with poor prognosis in serous ovarian cancer patients and is associated with therapeutic resistance

Analysis of publicly available datasets reveal that ERBB2 levels are elevated in several cancers including ovarian cancer^21^ (Fig. S1B). To understand how elevated ERBB2 levels impact prognosis in ovarian cancer patients, we analyzed survival data from both serous and endometrioid ovarian cancer patients. Patients were stratified according to ERBB2 expression levels in their tumors. While higher ERBB2 expression is associated with a shorter progression free survival in serous ovarian cancer patients, ERBB2 levels did not impact the progression free survival in patients with endometrioid ovarian cancers^22^ (Fig. 1I and S1C). As majority of high grade serous ovarian cancers are associated with p53 mutations, we analyzed if ERBB2 expression impacted prognosis in ovarian cancer patients who have p53 mutations in their tumor. As expected, high ERBB2 was associated with significantly worse prognosis in mutant p53 expressing ovarian tumors^22^ (Figs. 1J and S1D). Due to existing challenges with early detection, the majority of ovarian cancer patients continue to be diagnosed with metastatic disease^18^. Previous reports establish ZNF217 as a major driver of ovarian cancer metastasis^4^. Interestingly, ERBB2 expression is also highest in metastatic ovarian tumors^21^ (Fig. 1K). To understand how elevated ERBB2 levels in metastatic ovarian tumors correlates with survival outcome, we stratified patients with stage 3 and stage 4 ovarian tumors based on their ERBB2 levels. Remarkably, even in patients with advanced metastatic disease, elevated ERBB2 levels were associated with a worse prognosis (Fig. 1L).

A major clinical challenge in metastatic ovarian cancer is the emergence of chemoresistance^1–3^. Therefore, we sought to understand if ERBB2 expression in ovarian tumors can impact the clinical response to different chemotherapeutics that are currently used to treat ovarian cancer patients. Patients with high ERBB2 expression in their tumors responded poorly when treated with docetaxel, and topotecan (Figs. 1M and 1N). Similarly, the progression free survival, overall survival, and post-progression survival in ovarian cancer patients treated with paclitaxel was significantly worse in the high ERBB2 expressing cohort (Fig. 1O). Together, these data reveal that ERBB2 overexpression is associated with advanced metastatic disease and likely impacts the clinical outcome in these patients.

### ZNF217 overexpressing cells are more resistant to ERBB2-targeting therapeutics

ERBB2 levels are used as a predictive biomarker to identify patients who will benefit from anti-ERBB2 therapeutics^23–25^. As ZNF217 overexpression causes a significant increase in ERBB2 levels, we hypothesized that ovarian cancer cells with elevated ZNF217 levels will be more sensitive to ERBB2-targeting drugs. Confocal microscopy of non-permeabilized ovarian cancer cells confirmed higher ERBB2 levels on the cell surface in ZNF217 overexpressing OVCA420 cells (Fig. 2A). Trastuzumab is a humanized monoclonal antibody that is used to treat HER2/ERBB2 positive breast and stomach cancers^26, 27^. Mechanistically, trastuzumab binds to three distinct regions of domain IV in the extracellular domain of ERBB2 and inhibits downstream signaling^28^. Surprisingly, despite elevated ERBB2 levels, ZNF217 overexpressing OVCA420 cells were more resistant to trastuzumab (Figs. 2B and S2A). ERBB2 can undergo proteolytic cleavage to release a 90-95 kDa C-terminal proteolytic fragment^29^. This cytosolic truncated form of ERBB2 fragment is observed in 30% of ERBB2-positive breast cancers and has been implicated in trastuzumab resistance^29, 30^. Notably, confocal microscopy using permeabilized ovarian cancer cells revealed increased cytoplasmic ERBB2 signal in ZNF217 overexpressing OVCA420 cells (Fig. 2C). This was further confirmed by western blot analysis using control and ZNF217 overexpressing OVCA420 and TYK-Nu cells (Fig. 2D). To determine if the increased resistance to trastuzumab in ZNF217 overexpressing ovarian cancer cells is a result of elevated p95 ERBB2 levels, we used a cell-permeable small molecule ERBB2 kinase inhibitor, CP724,714. CP724,714, a selective ERBB2 inhibitor that has been shown to have more than 640-fold selectivity against EGFR and other receptor tyrosine kinases such as INSR, IRG-1R, PDGFR, VEGFR2, ABL, Src, and c-Met^31^. As CP724,714 will block signaling via both the full-length membrane bound ERBB2 as well as the intracellular p95-ERBB2, we argued that the ZNF217 overexpressing cells will be more sensitive to this drug. Contrary to our expectation, OVCA420-ZNF217 cells were more resistant to CP724,714 as compared to control cells (Figs. 2E and S2B). To confirm these results, and to ascertain that ZNF217 overexpressing cells do not demonstrate an increased sensitivity to ERBB2-targeting strategies, we tested the effect of an ERBB2 targeting PROTAC in ZNF217 overexpressing ovarian cancer cells. Consistent with its reported effect, treatment of OVCA420 and OVCA420-ZNF217 cells with ERBB2-PROTAC resulted in a decrease in ERBB2 levels (Fig. 2F). However, ZNF217 overexpressing OVCA420 and TYK-Nu cells were more resistant to ERBB2-PROTAC compared to control cells (Figs. 2G, 2H, S2C, and S2D). These data collectively show that despite elevated ERBB2 levels, ZNF217 overexpressing ovarian cancer cells are resistant to ERBB2-targeting strategies.

**Figure 2.**
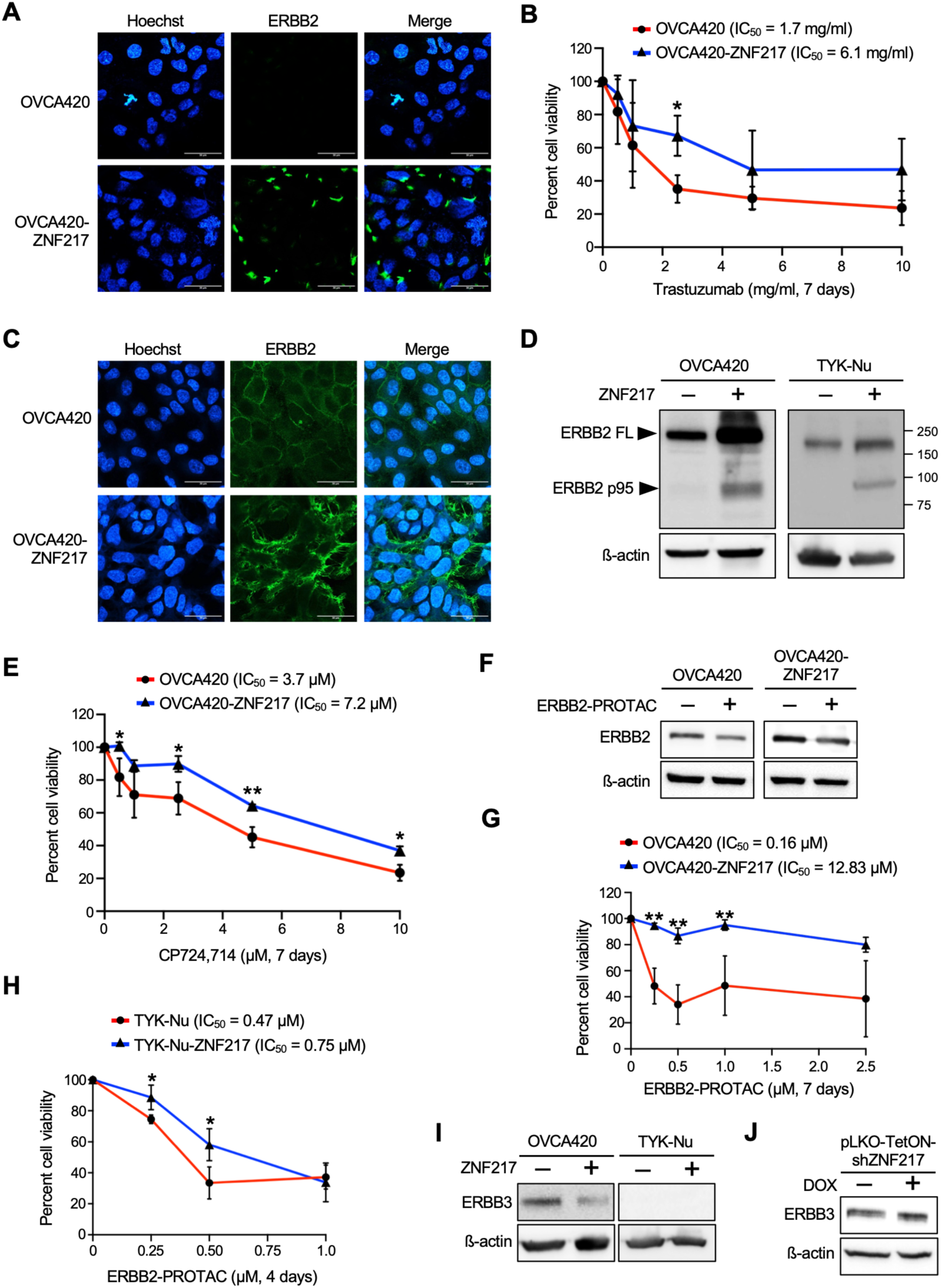
ZNF217-high ovarian cancer cells resist ERBB2-targeting therapeutics. **(A)** Confocal microscopy using non-permeabilized cells show elevated cell surface ERBB2 in OVCA420-ZNF217 cells. Blue – Hoechst (nucleus); Green – ERBB2. **(B)** Cell viability assay show OVCA420-ZNF217 cells are more resistant to trastuzumab (n=3, 7 days). **(C)** Confocal microscopy using permeabilized cells reveal cytoplasmic ERBB2 in OVCA420-ZNF217 cells. Blue – Hoechst (nucleus); Green – ERBB2. **(D)** Western blot showing elevated full-length and truncated p95 ERBB2 in OVCA420-ZNF217 and TYK-Nu-ZNF217 cells. **(E)** OVCA420-ZNF217 cells are more resistant to ERBB2 inhibitor CP724,714 (n=3, 7 days). **(F)** Western blot showing reduction in ERBB2 levels in OVCA420 and OVCA420-ZNF217 cells upon treated with ERBB2-PROTAC (2.5 µM, 24 h). **(G)** OVCA420-ZNF217 cells are more resistant to ERBB2-PROTAC (n=3, 7 days). **(H)** TYK-Nu-ZNF217 cells are more resistant to ERBB2-PROTAC (n=3, 4 days). **(I)** Western blot show that ZNF217 overexpression causes a decrease in ERBB3 levels in OVCA420 cells. No ERBB3 expression was detected in TYK-Nu and TYK-Nu-ZNF217 cells. **(J)** ZNF217 depletion increases ERBB3 levels in OVCA420 cells.

### ZNF217 promotes lapatinib resistance despite ERBB2 and EGFR upregulation

ERBB2 can form heterodimers with other receptor tyrosine kinases such as ERBB3 and EGFR (or ERBB1)^8, 9^. These heterodimers retain their ability to activate downstream signaling even when ERBB2 is blocked, allowing the cells to resist anti-ERBB2 targeting therapies^32^. For instance, as a heterodimer with ERBB2, ERBB3 can still bind to its ligand (neuregulin or heregulin) and activate the PI3K-AKT pathway even when ERBB2 is inhibited^32–35^. Therefore, we investigated if resistance to ERBB2-targeting therapeutics in ZNF217 overexpressing ovarian cancer cells is mediated by ERBB2’s ability to heterodimerize with either ERBB3 or EGFR. Contrary to the effect on ERBB2, ZNF217 overexpression resulted in a decrease in ERBB3 protein levels in OVCA420 cells (Fig. 2I). Consistently, ZNF217 knockdown led to a slight increase in ERBB3 levels in these cells (Fig. 2J). Interestingly, in both TYK-Nu and TYK-Nu-ZNF217 cells we did not see any detectable levels of ERBB3 (Fig. 2I). These data suggested that ERBB3 is unlikely to play a key role in the ability of ZNF217 overexpressing ovarian cancer cells to resist anti-ERBB2 therapeutics. Notably, ZNF217 was previously shown to positively regulate ERBB3 in breast cancer cells^36^. These differences between ovarian and breast cancer cells imply that the mechanisms by which ZNF217 impacts downstream targets and tumor progression can vary depending on the cancer cell type.

ERBB2 can also interact with EGFR to form heterodimers^9^. Western blot analysis showed that EGFR expression was increased in both OVCA420 and TYK-Nu cells following ZNF217 overexpression (Fig. 3A). Simultaneous inhibition of ERBB2 and EGFR can be achieved using lapatinib, a clinically approved oral tyrosine kinase inhibitor that is used to treat ERBB2-positive metastatic breast and esophageal cancers^37–39^. Remarkably, despite elevated ERBB2 and EGFR levels, the OVCA420-ZNF217 cells were more resistant to both short-term (72 hours) as well as long term (7 days) lapatinib treatment (Figs. 3B, 3C, 3D, and S3A). Similar results were also obtained using TYK-Nu-ZNF217 cells (Fig. 3E and S3B). These results suggest that while ZNF217 overexpressing ovarian cancer cells upregulate ERBB2 and EGFR, they are less reliant on these receptor tyrosine kinases for survival. Phosphorylation at Tyr877 of ERBB2 serves as a marker for ERBB2 activation in cancer cells^40^. Consistent with the observed phenotypes, despite elevated ERBB2 levels, ERBB2 phosphorylation at Tyr877 was substantially reduced in ZNF217 overexpressing OVCA420 and TYK-Nu cells, suggesting a loss of ERBB2 receptor kinase activity (Fig. 3F). These surprising results highlight the risk of using ERBB2 and EGFR levels as biomarkers to identify patients who are likely to respond to lapatinib without understanding the underlying downstream mechanism.

**Figure 3.**
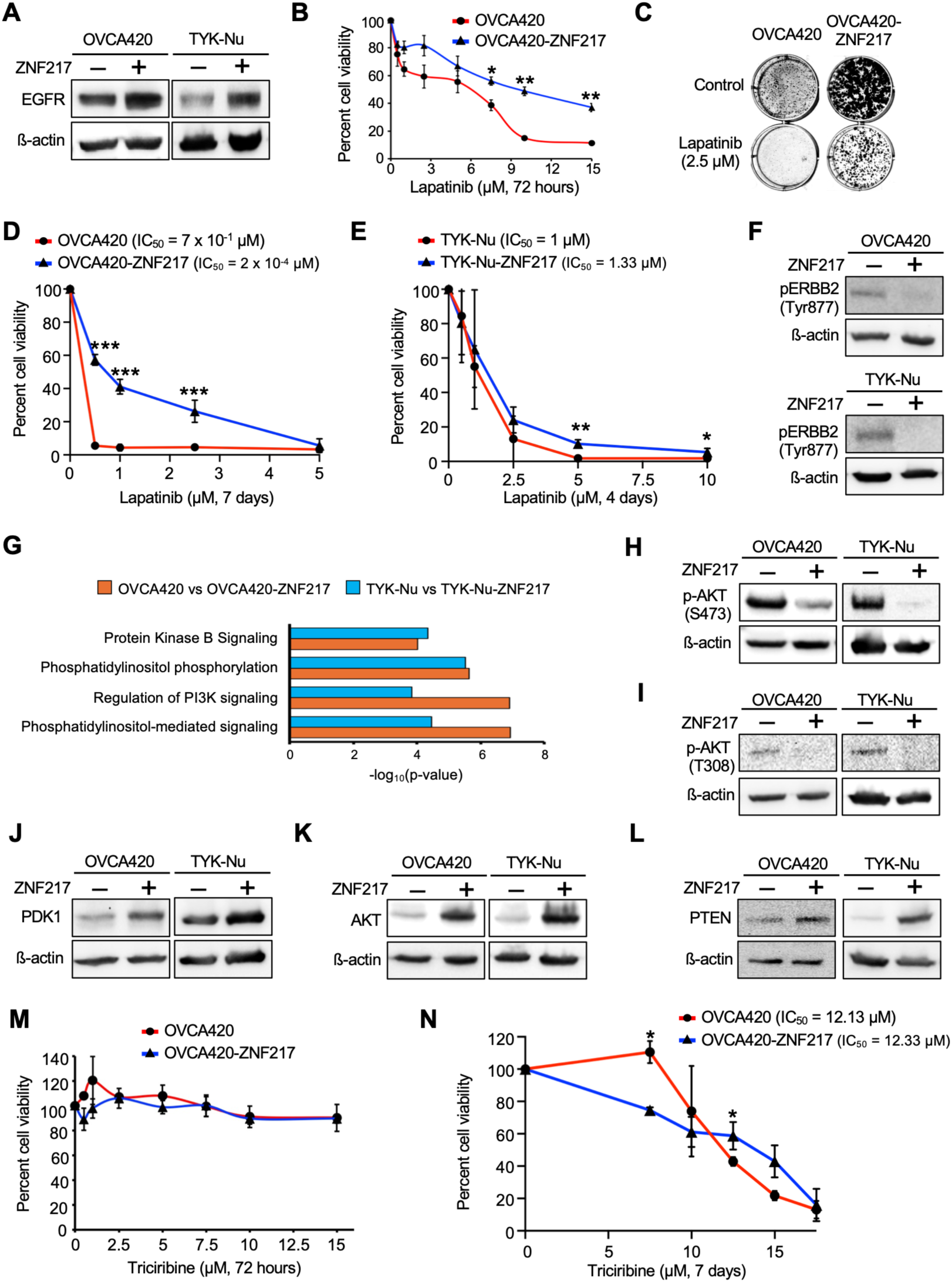
ZNF217-mediated resistance to anti-ERBB2 therapeutics is not dependent on EGFR and AKT signaling pathway. **(A)** Western blot showing that ZNF217 overexpression in OVCA420 and TYK-Nu cells results in upregulation of EGFR. **(B)** WST-1 cell viability assay show OVCA420-ZNF217 cells are more resistant to lapatinib (n=3, 72 h). **(C)** Representative Crystal violet staining showing increased resistance of OVCA420-ZNF217 cells to lapatinib (2.5 µM, 7 days). **(D)** OVCA420-ZNF217 cells are more resistant to long-term lapatinib treatment (n=3, 7 days). **(E)** TYK-Nu-ZNF217 cells are more resistant to lapatinib (n=3, 4 days). **(F)** Western blot showing that phosphorylation of Tyr877 in ERBB2 is reduced in ZNF217 overexpressing OVCA420 and TYK-Nu cells. **(G)** Protein Kinase B signaling related Gene Ontology terms are significantly enriched in the differentially expressed gene sets in ZNF217 overexpressing OVCA420 and TYK-Nu cells. **(H)** AKT Ser473 phosphorylation is reduced in ZNF217 overexpressing OVCA420 and TYK-Nu cells. **(I)** AKT Thr308 phosphorylation is reduced in ZNF217 overexpressing OVCA420 and TYK-Nu cells. **(J)** PDK1 levels are elevated in ZNF217 overexpressing OVCA420 and TYK-Nu cells. **(K)** AKT levels are elevated in ZNF217 overexpressing OVCA420 and TYK-Nu cells. **(L)** ZNF217 upregulates PTEN in OVCA420 and TYK-Nu cells. **(M)** WST-1 cell viability assay showing that both OVCA420 and OVCA420-ZNF217 cells are resistant to short-term triciribine treatment (n=3, 72 h). **(N)** Sensitivity of OVCA420 cells to long-term triciribine treatment (7 days) is not impacted by ZNF217 overexpression (n=3).

### ZNF217 overexpression results in decreased AKT activation in ovarian cancer cells

Alternative mechanisms that activate the PI3K-AKT pathway can bypass the inhibited ERBB2 and EGFR and drive resistance to drugs targeting these receptors^14, 41^. Therefore, we sought to determine the status of the AKT signaling pathway in ovarian cancer cells upon ZNF217 overexpression. RNAseq analysis in OVCA420 and TYK-Nu cells suggested that AKT signaling pathway is altered in these cells upon ZNF217 overexpression (Fig. 3G). Similar results were obtained using RNAseq data from OVCA420 cells in which ZNF217 was knocked down (Fig. S3C). Western blot analysis shows that phosphorylation of AKT at both Ser473 and Thr308 was reduced in ZNF217 overexpressing ovarian cancer cells (Figs. 3H and 3I). Phosphorylation at Ser473 and Thr308 are associated with AKT activation^42^. Therefore, these results suggest that the AKT signaling pathway is downregulated in ZNF217 overexpressing ovarian cancer cells. As commonly observed in cancer cells attempting to compensate for low-level or inhibited AKT signaling, we saw PDK1 and AKT is elevated at the protein level in ZNF217-high OVCA420 and TYK-Nu cells (Figs. 3J and 3K). RT-qPCR analysis using control and ZNF217 overexpressing OVCA420 cells revealed that the effect of ZNF217 on AKT abundance is not transcriptional (Fig. S3D). Interestingly, ZNF217 overexpression resulted in an increase in PTEN levels in both OVCA420 and TYK-Nu cells (Fig. 3L). Elevated PTEN levels can contribute towards reduced AKT activation observed in these cells and explains why AKT signaling maybe downregulated in ZNF217 overexpressing ovarian cancer cells despite elevated levels of many other pathway components. These results are different from previous findings in breast cancer cells, where ZNF217 overexpression was reported to activate the PI3K–AKT signaling pathway, potentially through an ERBB3-dependent mechanism^43, 44^. Consistent with activation of AKT in breast cancer cells, the ZNF217 overexpression sensitized these cells to the allosteric AKT inhibitor, triciribine^43, 44^. However, in ovarian cancer cells, ZNF217 overexpression did not alter sensitivity to triciribine under either short-term (72 h) or long-term (7 day) treatment conditions (Figs. 3M, 3N, and S3E). Again, these results suggest that the mechanisms that ZNF217 co-opts to promote oncogenesis and therapeutic resistance can vary between cancer cell types and depending on the context. Therefore, therapeutic approaches in ZNF217 overexpressing tumors should carefully consider the underlying mechanisms that are critical in sustaining tumor growth and chemoresistance.

### ZNF217 activates AXL to promote oncogenic phenotypes in ovarian cancer cells

In addition to interacting with ERBB3 and EGFR, ERBB2 can form heterodimers with other receptor tyrosine kinases such as the fibroblast growth factor receptors and AXL^10–12^. Resistance to ERBB2-targeted therapies can arise due to heterodimerization of ERBB2 with these receptors as well as through ERBB2-independent signaling mediated by these receptor tyrosine kinases^10–12^. Therefore, we investigated the effect of ZNF217 on both fibroblast growth factor receptors and AXL in ovarian cancer cells. Remarkably, ZNF217 overexpression resulted in an increase in FGFR1, FGFR4, and AXL levels in both OVCA420 and TYK-Nu cells (Figs. 4A, 4B, and 4C). Therefore, ZNF217 upregulates several receptor tyrosine kinases in ovarian cancer cells including ERBB2, EGFR, FGFR1, FGFR4, and AXL, thereby allowing these cells to respond rapidly to different growth factors in the tumor microenvironment. Further, by establishing signaling plasticity, ZNF217 enables these cells to bypass growth inhibitory signals such as those induced by chemotherapeutic agents. Among the receptor tyrosine kinases examined, ZNF217 overexpression had the most pronounced effect on AXL levels, aside from ERBB2 (Fig. 4C). RT-qPCR analysis shows that ZNF217 overexpression in OVCA420 cells results in significant upregulation of AXL mRNA, suggesting the effect to be transcriptional (Fig. 4D). RNAseq analysis reveals that in addition to AXL, its ligand GAS6, and dimerization partner EGFR is upregulated upon ZNF217 overexpression (Fig. 4E). ZNF217 overexpression also caused downregulation of another ligand PROS1 suggesting that GAS6 is likely to be the major ligand for AXL in this context (Fig. 4E). Consistent with these data, ZNF217 overexpression increased phosphorylation of AXL at Tyr779, indicating enhanced AXL signaling activity in these cells (Fig. 4F). Clinical data reveal that elevated AXL levels in ovarian tumors are associated with shorter overall and progression free survival^22^ (Figs. 4G and 4H). AXL overexpression is also associated with poor response to platin- and taxol-based chemotherapeutics as well as gemcitabine^22^ (Figs. S4A - S4H). As AXL activity is elevated in ZNF217 overexpressing cells, we assessed whether ZNF217 overexpression sensitizes ovarian cancer cells to AXL inhibition. Compared to control cells, both OVCA420-ZNF217 and TYK-Nu-ZNF217 cells exhibited greater decrease in cell viability when treated with the AXL inhibitor TP-0903 (Figs. 4I, 4J, and S4I). As ZNF217 enhances the ability of ovarian cancer cells to migrate, we tested if AXL inhibition can reverse this phenotype^4^. Transwell migration assay reveal that TP-0903 treatment abolished the migratory advantage ZNF217 confers to ovarian cancer cells (Figs. 4K and 4L). These results show that AXL is an important downstream effector of ZNF217’s oncogenic function in ovarian cancer cells. Further, our data also suggest that AXL has potential as an actionable therapeutic target in ZNF217-driven ovarian tumors.

**Figure 4.**
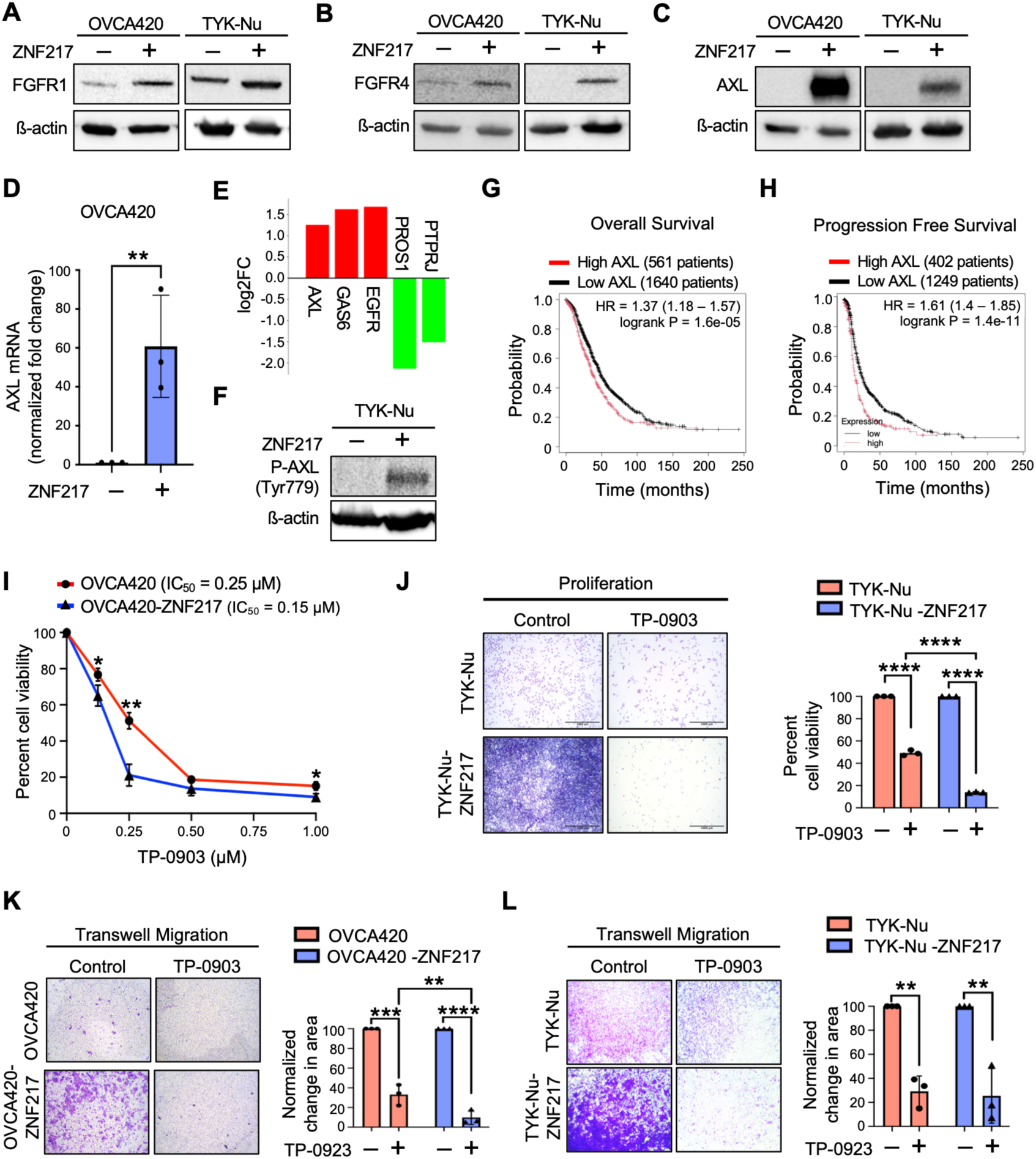
ZNF217 activates AXL to promote oncogenic phenotypes in ovarian cancer cells. **(A)** Western blot showing upregulation of FGFR1 in OVCA420-ZNF217 and TYK-Nu-ZNF217 cells. **(B)** Western blot showing upregulation of FGFR4 in OVCA420-ZNF217 and TYK-Nu-ZNF217 cells. **(C)** Western blot showing upregulation of AXL in OVCA420-ZNF217 and TYK-Nu-ZNF217 cells. **(D)** RT-qPCR data show that ZNF217 overexpression in OVCA420 cells causes an increase in AXL mRNA levels. Values were normalized to TBP mRNA. **(E)** RNAseq data reveal that genes associated with AXL activation are altered upon ZNF217 overexpression in TYK-Nu cells. **(F)** AXL Tyr779 phosphorylation is elevated in TYK-Nu-ZNF217 cells. **(G)** Ovarian Cancer patients with AXL-high tumors have lower overall survival. Data retrieved from Kmplotter. **(H)** Ovarian cancer patients with AXL-high tumors have lower progression free survival. Data retrieved from Kmplotter. **(I)** Cell viability assay shows that OVCA420-ZNF217 cells are more sensitive to the AXL inhibitor TP-0903 (n=3, 7 days). **(J)** Cell viability assay shows that TYK-Nu-ZNF217 cells are more sensitive to the AXL inhibitor TP-0903 (n=3, 2.5 µM, 24 h). **(K)** Transwell assay shows that TP-0903 (100 nM) reverses ZNF217 induced migratory phenotype in OVCA420 cells. **(L)** Transwell assay shows that TP-0903 (100 nM) reverses ZNF217 induced migratory phenotype in TYK-Nu cells.

### MAPK signaling is critical for ZNF217 driven metastatic phenotypes in ovarian cancer cells

Receptor tyrosine kinases such as AXL, EGFR, FGFRs as well as ERBB2 are known to activate MAPK kinases^45–47^. Therefore, we investigated the status of MAPK signaling pathway in ZNF217 overexpressing ovarian cancer cells. RNAseq analysis reveals that ZNF217 overexpression causes activation of the ERK/MAPK signaling cascade in both OVCA420 and TYK-Nu cells (Figs. 5A and 5B). Similar results were obtained using RNAseq data from control and ZNF217 knockdown OVCA420 cells (Fig. S5A). Next, we investigated if ZNF217 overexpression impacted ERK abundance in ovarian cancer cells. RT-qPCR analysis revealed that ZNF217 overexpression did not alter ERK mRNA levels in OVCA420 cells (Fig. 5C). Interestingly, ZNF217 overexpression in both OVCA420 and TYK-Nu cells resulted in a dramatic increase in ERK1/2 levels, implying ZNF217 upregulates ERK levels in these cells via post-transcriptional mechanisms (Fig. 5D). These data also suggested that ZNF217 overexpressing ovarian cancer cells are primed to respond to upstream activating signals that activate MAPK signaling. Consistent with active signaling via the MAPK pathway, phosphorylation of ERK1/2 at Thr202/Tyr204 is elevated in ZNF217 overexpressing OVCA420 and TYK-Nu cells (Fig. 5E). Next, we tested if active ERK signaling is critical for ZNF217 to promote pro-metastatic phenotypes such as cell migration and invasion in ovarian cancer cells. Treatment of OVCA420 with ERK inhibitor LY3214996 induced a profound decrease in cell migration in ZNF217 overexpressing cells but not control cells (Fig. 5F). Similarly, LY3214996 also abolished ZNF217’s ability to promote invasion in ovarian cancer cells through Matrigel (Fig. 5G). These data establish activation of MAPK signaling as a key downstream mechanism that mediates ZNF217’s oncogenic function in ovarian cancer cells (Fig. 5G). As AXL is known to activate ERK signaling, we evaluated the combined effect of co-targeting AXL and ERK on ovarian cancer cell migration and invasion. While co-treating ZNF217 overexpressing ovarian cancer cells with TP-0903 (AXL inhibitor) and LY3214996 (ERK inhibitor) resulted in a profound decrease in cell migration (Fig. S5B) and invasion (Fig. S5C), the effect was not significantly more than targeting either of these kinases alone (Figs. 4K, 4L, 5F and 5G), suggesting that the effect of AXL on these phenotypes is likely mediated via ERK activation.

**Figure 5.**
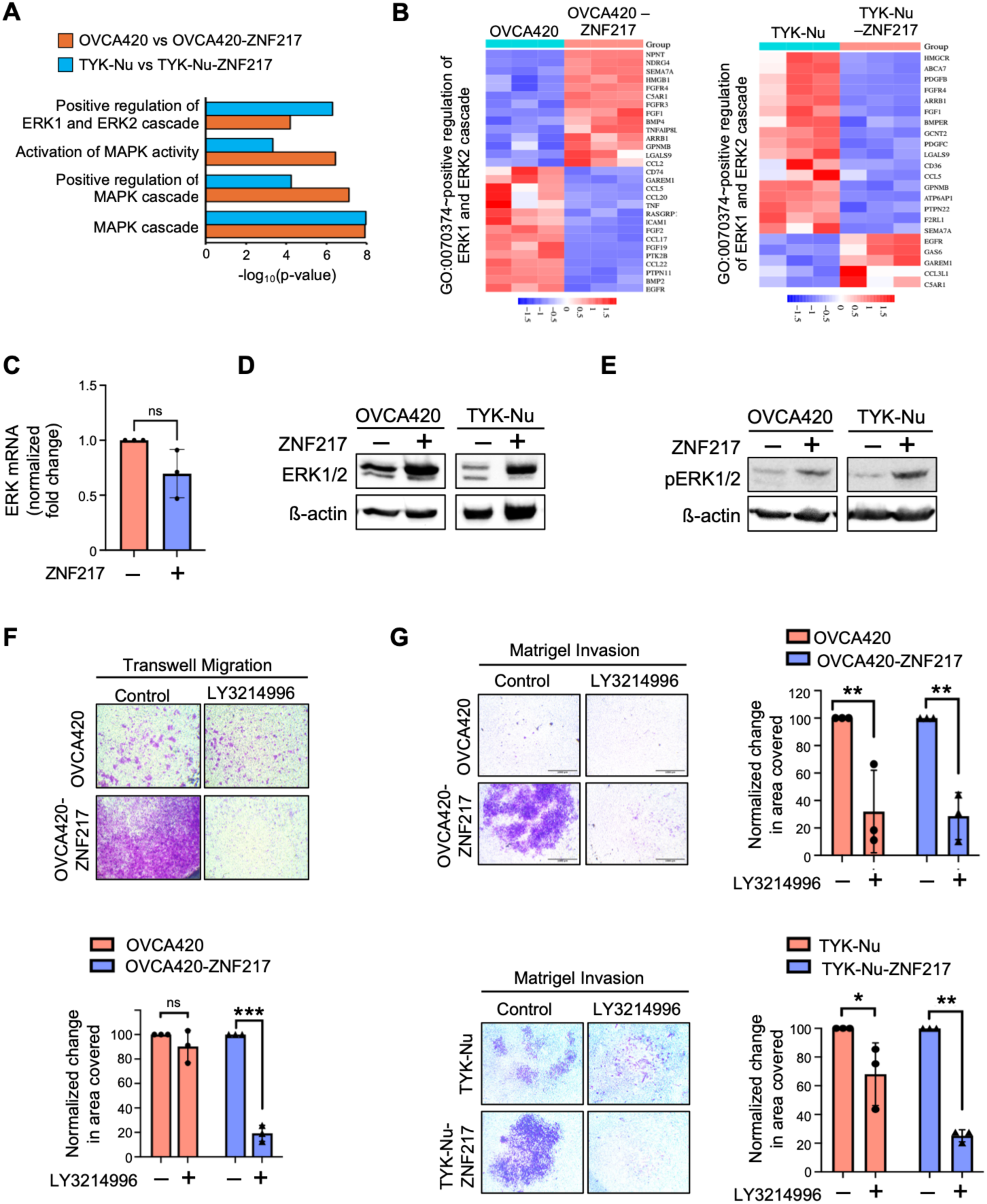
MAPK signaling is critical for ZNF217 to drive metastatic phenotypes in ovarian cancer cells. **(A)** RNAseq data reveal that MAPK signaling related Gene Ontology terms are significantly enriched upon ZNF217 overexpression in OVCA420 and TYK-Nu cells. **(B)** Heatmap showing genes related to ERK1/2 signaling that are differentially regulated in OVCA420-ZNF217 and TYK-Nu-ZNF217 cells. **(C)** RT-qPCR data reveal that ZNF217 overexpression does not alter ERK mRNA levels in OVCA420 cells. **(D)** Western blot showing elevated ERK1/2 levels in OVCA420-ZNF217 and TYK-Nu-ZNF217 cells. **(E)** Western blot reveal elevated phospho-ERK1/2 levels in ZNF217 overexpressing OVCA420 and TYK-Nu cells. **(F)** Transwell assay shows that ERK inhibitor LY3214996 (1 µM) decreases cell migration in OVCA420-ZNF217 cells. **(G)** LY3214996 (1 µM) reverses ZNF217 induced invasion in OVCA420 and TYK-Nu cells in Matrigel invasion assay.

### AXL activation drives MAPK signaling in ZNF217-overexpressing ovarian cancer cells

To determine whether upregulation and activation of AXL in ZNF217-overexpressing cells promotes MAPK signaling, we examined the effects of the AXL inhibitor TP-0903 on ERK expression and activation in TYK-Nu-ZNF217 and OVCA420-ZNF217 cells. Western blot analysis showed that TP-0903 treatment did not alter total ERK protein levels in either cell line (Fig. 6A). However, TP-0903 induced a profound decrease in ERK phosphorylation in both OVCA420-ZNF217 and TYK-Nu-ZNF217 cells (Fig. 6B). These findings suggest that ZNF217-mediated AXL upregulation and activation is a key driver of MAPK pathway activation in these cells. Next, we examined whether AXL activation affects ERBB2 and AKT expression and activity in ZNF217-overexpressing ovarian cancer cells. AXL inhibition did not alter total ERBB2 protein levels (Fig. 6C). However, TP-0903 treatment decreased ERBB2 phosphorylation at Tyr877, suggesting that AXL may positively regulate ERBB2 phosphorylation and ERBB2-mediated signaling in these cells (Fig. 6D). In contrast, AXL inhibition did not alter total AKT levels (Fig. 6E) or AKT phosphorylation (Fig. 6F). AXL overexpression has been previously reported to be associated with acquired resistance to anti-ERBB2 treatment in ERBB2-positive breast cancer^11^. Further, in ERBB2-amplified breast cancer patients with high AXL expression, combining AXL inhibition with trastuzumab was shown to be an effective strategy^11^. Therefore, we investigated if combining trastuzumab with AXL inhibition, either alone or in combination with ERK inhibition, enhances the cytotoxic effects on ZNF217-overexpressing ovarian cancer cells. Consistent with a central role for AXL–ERK signaling in the survival of ZNF217-overexpressing ovarian cancer cells, AXL inhibition, either alone or in combination with an ERK inhibitor, resulted in a profound reduction in cell viability (Figs. 6G and 6H). Notably, combining TP-0903 with trastuzumab produced a further significant decrease in cell viability (Fig. 6G). Similar effects were observed when trastuzumab was combined with both AXL and ERK inhibitors (Fig. 6H). These findings suggest that ZNF217-overexpressing cells that persist following AXL inhibition may increasingly engage alternative receptor tyrosine kinase signaling pathways, including ERBB2, to sustain cell survival, potentially rendering these cells more susceptible to therapeutic strategies targeting these pathways.

**Figure 6.**
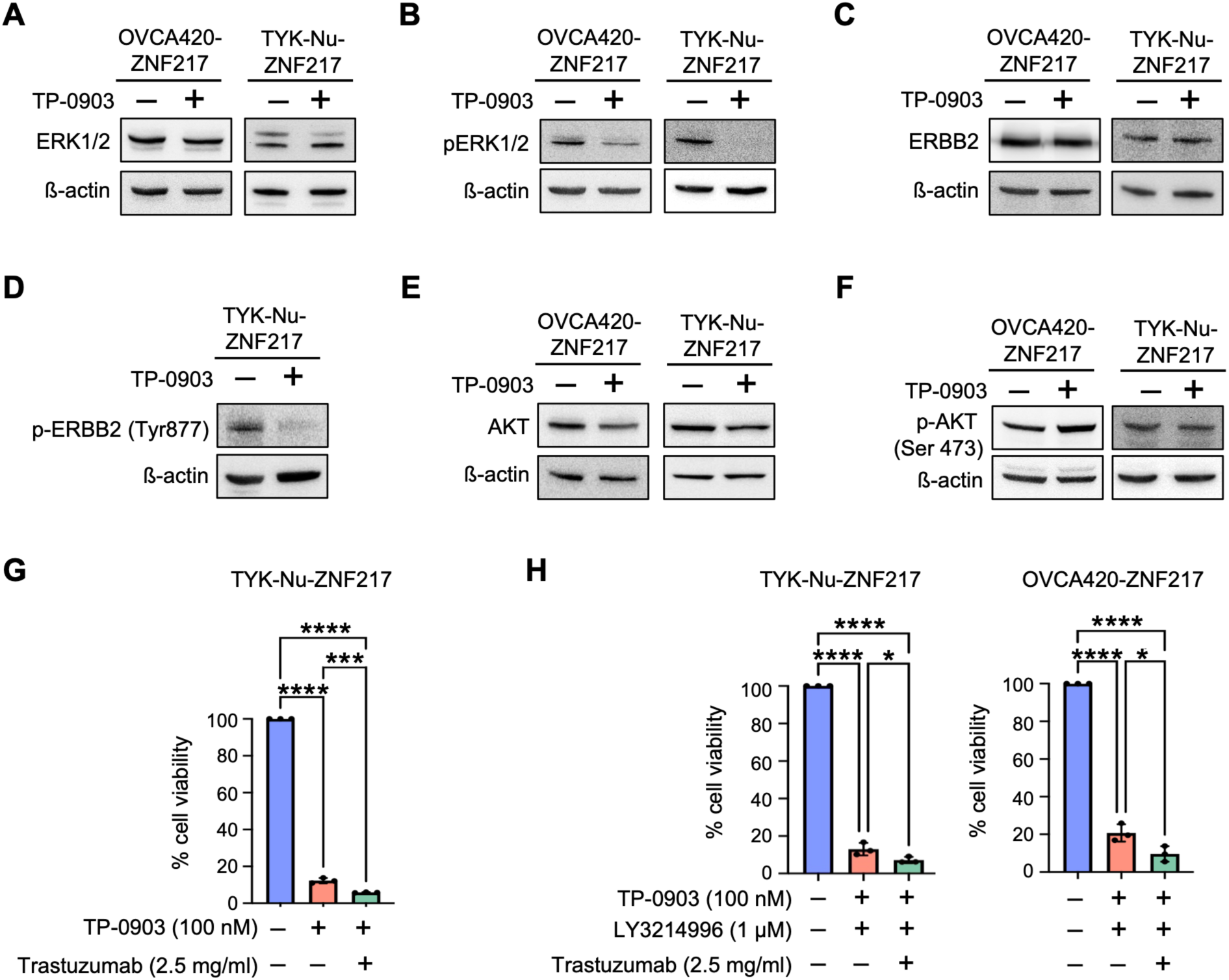
AXL activation drives MAPK signaling in ZNF217-overexpressing ovarian cancer cells. **(A)** TP-0903 treatment (100 nM, 24h) does not alter the total ERK protein level in ZNF217 overexpressing OVCA420 and TYK-Nu cells. **(B)** TP-0903 treatment (100 nM, 24h) decreases ERK phosphorylation in ZNF217 overexpressing OVCA420 and TYK-Nu cells. **(C)** TP-0903 treatment (100 nM, 24h) does not alter the total ERBB2 protein level in ZNF217 overexpressing OVCA420 and TYK-Nu cells. **(D)** TP-0903 treatment (100 nM, 24h) decreases ERBB2 phosphorylation (Tyr 877) in TYK-Nu-ZNF217 cells. TP-0903 treatment (100 nM, 24h) does not alter **(E)** total AKT and **(F)** phosphor-AKT (Ser473) levels in ZNF217 overexpressing OVCA420 and TYK-Nu cells. **(G)** Trastuzumab (2.5 mg/ml) further decreases cell viability beyond the effect of TP-0903 (100 nM) in TYK-Nu-ZNF217 cells (7 days treatment). **(H)** Trastuzumab (2.5 mg/mL) further decreases cell viability beyond the effect of combined TP-0903 (100 nM) and LY3214996 (1 µM) treatment in ZNF217-overexpressing ovarian cancer cells (7 days treatment).

### AXL inhibition impairs ovarian tumor progression and improves survival in vivo

Given that AXL is upregulated in ZNF217-overexpressing ovarian cancer cells and plays an important role in mediating the oncogenic effects of ZNF217, we sought to determine whether ZNF217 and AXL expression are correlated in human ovarian tumors. To this end, we analyzed publicly available datasets, including TCGA and GTEx, using the GEPIA (Gene Expression Profiling Interactive Analysis) platform to assess the relationship between ZNF217, AXL, and ERBB2 expression in ovarian tumors^48^. Our analysis revealed a statistically significant positive correlation between ZNF217 and AXL expression, as well as between ZNF217 and ERBB2 expression, in ovarian tumors (Fig. 7A). To determine whether AXL is critical for ZNF217-driven ovarian tumor progression and to evaluate its potential as a therapeutic target in ZNF217-overexpressing ovarian tumors, we investigated the effects of the AXL inhibitor TP-0903 on tumor burden and survival using a mouse intraperitoneal xenograft model. Consistent with our in vitro findings, TP-0903 treatment reduced metastatic tumor burden (Figs. 7B and 7C) and significantly increased survival in a dose-dependent manner (Fig. 7D). These findings provide in vivo evidence that AXL inhibition may represent a potential therapeutic strategy for targeting ZNF217-overexpressing ovarian tumors.

**Figure 7.**
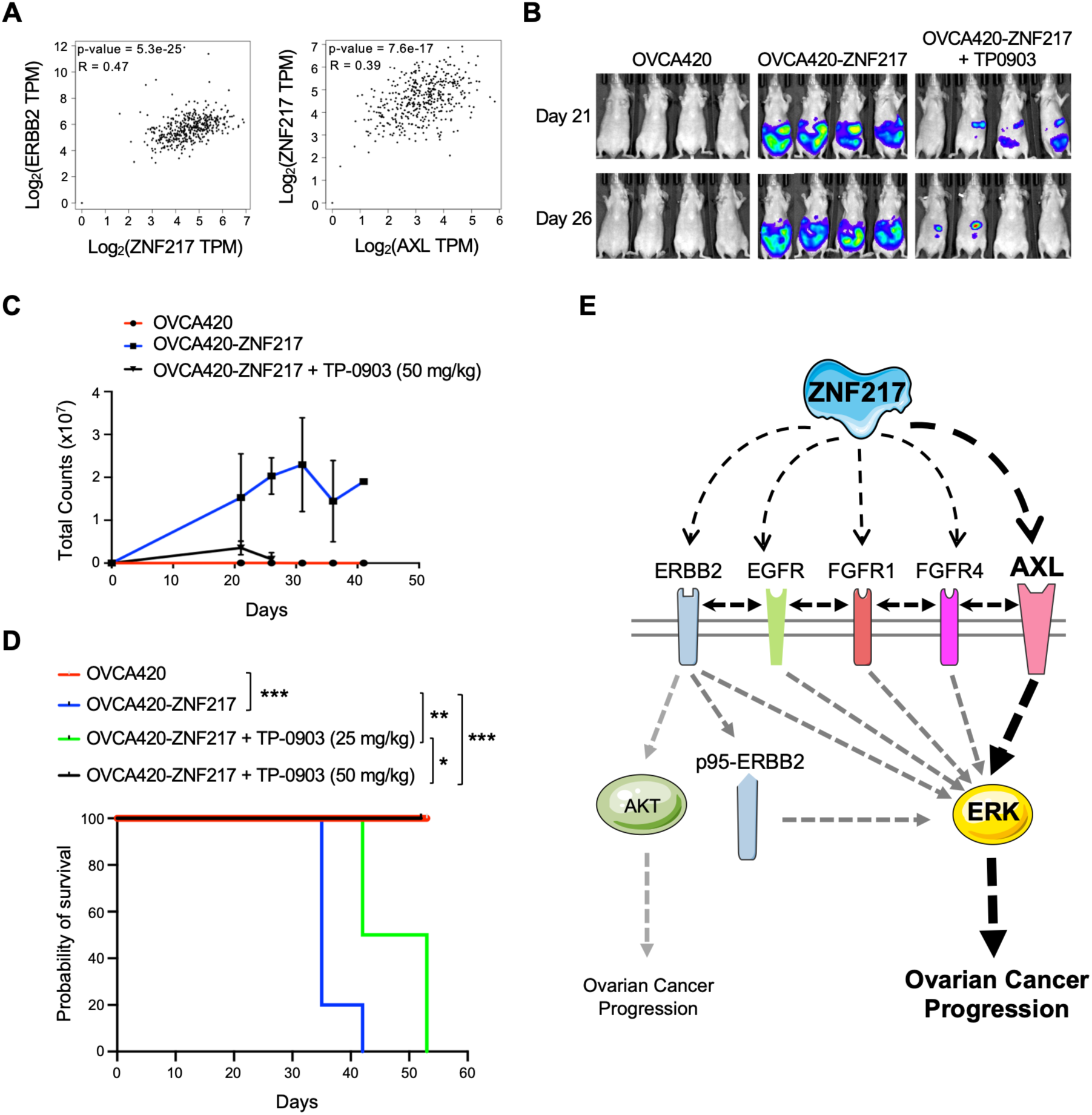
AXL inhibition suppresses ZNF217-driven ovarian tumor growth and improves survival in vivo. **(A)** Correlation analysis using data retrieved from the GEPIA (Gene Expression Profiling Interactive Analysis) platform show positive correlation between ZNF217 and ERBB2 (left) and ZNF217 and AXL (right) in ovarian tumors. R value shown is Spearman correlation coefficient. **(B)** Representative bioluminescence images from female Foxn1 mice i.p. injected with luciferase-tagged OVCA420 and OVCA420-ZNF217 cells reveal significant reduction in ZNF217-driven tumor burden upon TP-0903 treatment (50 mg/ml). **(C)** Quantification of in vivo imaging in Foxn1 nude mice show that AXL inhibitor (TP-0903, 50 mg/kg) reduces progression of ZNF217-driven metastatic ovarian tumor in mice (n = 6 mice/group). **(D)** Kaplan-Meier curve reveals that TP-0903 treatment increases survival in mice bearing ZNF217-overexpressing ovarian tumors in a dose dependent manner (n=6 mice/group). **(E)** Model describing key findings reported in this study.

Taken together, the data presented in this manuscript demonstrate that ZNF217 upregulates several receptor tyrosine kinases in ovarian cancer cells, thereby creating multiple potential signaling pathways through which these cells can respond to changes in growth-promoting and inhibitory signals (Fig. 7E). The broad repertoire of receptor tyrosine kinases, together with their ability to heterodimerize, homodimerize, or signal independently in a context- and cell type-dependent manner, may confer substantial signaling plasticity to ZNF217-overexpressing cancer cells. Being primed to rewire cellular signaling in response to external stimuli may contribute to their ability to adapt to and resist therapeutic interventions targeting individual receptor tyrosine kinases (Fig. 7E). Despite the availability of these potential alternative signaling pathways, our findings indicate that ZNF217-overexpressing ovarian cancer cells remain particularly dependent on the AXL–ERK signaling axis for maintenance of oncogenic phenotypes and cell survival. Thus, the AXL–ERK signaling axis represents a therapeutically actionable vulnerability in ZNF217-overexpressing ovarian cancer cells.

## DISCUSSION

Chemoresistance is a major clinical challenge that has limited the effectiveness of extant therapeutics in metastatic ovarian cancer. Overcoming this challenge will require identifying factors that promotes drug resistance, understanding the mechanistic basis of their action, and developing strategies to target them. Equally important to improve clinical outcome in ovarian cancer patients is discovering reliable biomarkers that can distinguish potential responders from non-responders. Upregulation of the oncogene ZNF217 was recently shown to enable ovarian cancer cells to resist chemotherapeutic drugs^4^. However, the underlying mechanisms through which ZNF217 drives these effects remained poorly defined. We show that ZNF217 upregulates multiple key receptor tyrosine kinases as well as components of the downstream MAPK signaling pathway that they activate (Fig. 5G). This confers these cells the ability to rewire signaling pathway in response to changes in external microenvironment including chemotherapeutic stress.

ERBB2 is among the receptor tyrosine kinases that is upregulated in ZNF217 overexpressing ovarian cancer cells. ERBB2 is a well-established oncogene that drives tumor progression and therapeutic resistance across several solid tumors^37^. Therapeutics that target ERBB2 are standard-of-care treatment of ERBB2-positive breast and gastric cancers^37^. Effective treatment strategies are often guided by reliable biomarkers that help distinguish potential responders from non-responders. ERBB2 levels are used to identify breast and gastric cancer tumors that are likely to respond to anti-ERBB2 drugs^16^. As ERBB2 levels are elevated in ZNF217 overexpressing tumors, we hypothesized that ZNF217-high ovarian tumors be sensitive to anti-ERBB2 therapeutics. However, we discovered that ZNF217 overexpressing cells were more resistant to multiple ERBB2 targeting drugs. These results suggest that ZNF217-high ovarian cancer cells either are not reliant on ERBB2 for survival and are able to rewire cellular signaling to resist ERBB2 inhibition. These results highlight the risk of using ERBB2 levels alone as a biomarker to identify patients who are potential responders without understanding the underlying mechanisms that drive tumor progression.

Multiple mechanisms are known to contribute to anti ERBB2-resistance in ERBB2-high tumors^32^. Heterodimerization with EGFR has been shown to enable cancer cells resist ERBB2 inhibition. However, ZNF217-high tumors were resistant to combined inhibition of ERBB2 and EGFR suggesting that these cells are reliant on other mechanisms. While inhibitors of EGFR and ERBB2 have been successfully in the treatment of multiple solid tumors, clinical trials using these compounds have been discouraging in ovarian cancer^49^. Signaling plasticity in ovarian cancer cells and activation of compensatory pathways are potential mechanisms that enable ovarian tumors to resist these drugs. As activation of AKT signaling pathway is an important downstream mechanism co-opted by ERBB2-high tumors, we investigated the status of AKT signaling in ZNF217 overexpressing ovarian cancer cells. Unlike in breast cancer cells, ZNF217 upregulation in ovarian cancer cells did not lead to AKT activation^43^. In fact, AKT signaling was downregulated in ZNF217-high ovarian cancer cells suggesting that these cells are reliant on alternate effectors to drive tumor progression. PI3K inhibition was shown to cause acquired ERK-dependency in ERBB2 overexpression breast cancer cells^50^. Consistently, we observed that the ZNF217-high ovarian cancer cells are dependent on MAPK signaling pathway to drive oncogenic phenotypes.

While multiple receptors that were upregulated in ZNF217-high ovarian cancer cells are capable of activating MAPK signaling, the effect of ZNF217 on AXL expression was particularly profound. AXL is a member of the TAM (TYRO3, AXL, MER) family of receptor tyrosine kinases that has been linked to cell viability, drug resistance, epithelial-to-mesenchymal transition, and metastasis in several different cancers^51–53^. Consistent with these, AXL inhibition reversed ZNF217 induced pro-metastatic phenotypes such as increased cell migration and invasion. AXL inhibition is being explored as a potential therapeutic strategy in multiple cancers including triple negative breast cancers^54–57^. Interestingly, AXL overexpression was shown to drive acquired resistance to ERBB2 targeting therapeutics in breast cancer cells^58^. AXL’s ability to heterodimerize with ERBB2 was shown to contribute to anti-ERBB2 therapeutics in breast cancer patients^11^. Consistently, AXL overexpression was shown to predict poor prognosis in ERBB2-positive breast cancer patients^11^. Despite being an important driver of oncogenic phenotypes, the mechanisms that mediates the frequent overexpression of AXL remains poorly understood^59^. In this report, we show that ZNF217 overexpression results in a profound increase in AXL levels in ovarian cancer cells. As ZNF217 upregulates ERBB2 in addition to AXL in ovarian cancer cells, heterodimerization of these receptors is likely to contribute to the observed resistance to anti-ERBB2 therapeutics observed in ZNF217-high ovarian cancer cells. AXL and its ligand Gas6 are upregulated in ovarian tumors, particularly in tumors that are platinum resistant^60^. Combining AXL targeting-nanobodies and olaparib exerted a synergistic anti-tumor effect in an orthotopic ovarian cancer mouse model^60^. Future studies that investigate mechanisms that regulate AXL levels in ovarian cancer cells and determine the effect of disrupting AXL signaling in ovarian tumors will be critical in translating AXL’s potential as a therapeutic target in chemoresistant ovarian tumors.

## METHODS

### Chemicals and Reagents

RPMI-1640 media containing L-glutamine was purchased from Corning (#10-040-CMR). Fetal Bovine Serum (FBS) was obtained from Omega scientific (#FB-02). Penicillin-streptomycin (Pen-Strep) was purchased from Gibco (#15140163). Cell Proliferation reagent WST-1 was purchased from Millipore-Sigma (# 11644807001). PBS was obtained from Corning (# 21-040-CV) and 4% paraformaldehyde was purchased from Thermofisher (#J19943). 8 µm, 24 well-permeable insert was purchased from Greiner Bio-One Thincert^TM^ (# 662638). Matrigel was purchased from Corning (# 354234). Cover glass (0.15mm thickness) was obtained from Carolina (# 633029). Prolong™ Gold Antifade mounting media was purchased from Thermofisher (# P36930). Trastuzumab (# 83967-1) was purchased from BPS Bioscience. ERBB2-PROTAC (# 7886) was obtained from TOCRIS. Lapatinib (# 11493), CP724,714 (# 19172), tricirbine (# 10010237), LY3214996 (# 27936), TP0903 (# 28757) were obtained from Cayman chemicals.

### Cell lines and culture condition

OVCA420 and TYK-Nu cells were a kind gift from Dr. Kwong-Kwok Wong (MD Anderson Cancer Center) and confirmed to be negative for mycoplasma. All human ovarian cancer cell lines were grown in RPMI-1640 containing L-glutamine supplemented with 10% FBS and 1% Pen-Strep. HEK293T cells were used to generate lentivirus particles. HEK293T cells were grown in DMEM media supplemented with 10% FBS and 1% Pen-Strep. All cell lines were grown in 37 °C incubator with 5% CO_2_. ZNF217 stable overexpression cell lines, OVCA420-RFP-ZNF217 and TYKNu-ZNF217 and appropriate control cell lines were generated using lentiviral transfection method as described previously^4^. To generate stable OVCA420 cells with doxycycline inducible ZNF217 knockdown, validated shRNA targeting ZNF217 (TGTAAAGTTCAAA) was cloned into the lentiviral vector pLKO.1-Tet-ON (Addgene #21915) using previously published protocol^61^. pLKO.1-Tet-ON containing ZNF217 shRNA was confirmed by sanger sequencing and used to generate lentivirus particles using HEK193T cells as described previously^4^. The lentivirus was transduced into OVCA420, and stably transfected cells were selected using puromycin and confirmed by western blot.

### Western blot analysis

Cells were lysed using RIPA buffer supplemented with protease and phosphatase inhibitors. Protein concentrations were determined using Bio-Rad Protein Assay Dye Reagent. Equal amount of total protein was run using an SDS-PAGE gel and transferred to a PVDF membrane. The membrane was blocked for 1 hour at room temperature in 2% milk, incubated with primary antibodies overnight at 4°C on a shaker. The membranes were washed three times with 1X TBST, incubated with appropriate HRP-linked secondary antibodies for 1 hour at room temperature on a shaker, washed again with 1X TBST three times, developed using enhanced chemiluminescence reagent, and imaged using a BioRad ChemiDoc Imager. The list of antibodies used in this study are provided in Table S1.

### Cell viability assays

For short-term drug viability assays, 5000 cells were seeded in each well of a 96-well plate. After 24 hours incubation at 37 °C with 5% CO_2_, the cells were treated with respective drugs and incubated for an additional 72 hours. Viable cells in each well were determined using WST-1 reagent as per the manufacturer’s recommendation. For long-term cell viability assays, 5000 cells were seeded in each well of a 6-well plate. After 24 hours incubation at 37 °C with 5% CO_2_, the cells were treated with different chemotherapeutic drugs. OVCA420 cells were incubated for an additional 7 days, and TYK-Nu cells were incubated for 4 days. Post-drug treatment, the cells were washed with 1x PBS, fixed with 4% Paraformaldehyde and stained using 0.5% Crystal Violet. Additional Crystal violet was removed by washing the wells with distilled water. Cells were imaged using microscope and quantified with Image J.

### Transwell cell migration assays

4 X 10^4^ control and ZNF217 overexpressing cells were resuspended in RPMI-1640 media containing 10% FBS and seeded inside 8 µM 24 well transwell inserts. The insert was placed in a well in a 24 well plate containing RPMI-1640 media with 20% FBS. Cells were incubated at 37 °C with 5% CO_2_ for 24 hours. At the end of 24 hours, inserts were washed with 1x PBS, fixed with 4% Paraformaldehyde and stained using 0.5% Crystal Violet. Additional Crystal violet was removed by washing the inserts with distilled water. The non-migrated cells were removed by wiping inside of inserts using cotton swab. Cells that successfully migrated through the transwell membrane were imaged using microscope and quantified with Image J.

### Matrigel cell invasion assays

Corning Matrigel^®^ Matrix was thawed overnight at 4 °C. The Matrigel was diluted at a ratio of 1:3 with serum free RPMI-1640 containing L-glutamine. 30 µl of diluted Matrigel was used to coat each 8 µm, 24 well-permeable insert and allowed to jellify for 30 minutes at room temperature. 4 X 10^4^ cells (TYK-Nu) and 8 X 10^4^ cells (OVCA420) in media containing no serum were seeded inside the Matrigel coated inserts. The inserts were places in a well containing RPMI-1640 + L-glutamine containing 20% Fetal Bovine serum and 1% penicillin and streptomycin. The cells were incubated for 24 hours and 48 hours for TYK-Nu and OVCA420 respectively. Subsequently, they were washed with 1X PBS, fixed with 4% paraformaldehyde, stained with 0.5 % Crystal violet, and quantified using Image J as mentioned in transwell migration protocol.

### Immunofluorescence and Confocal Microscopy

Cover glass (0.15mm thickness) was coated with 0.05 mg/ml poly-D lysine for 1 hour at 37 °C and washed with distilled water and left to dry before seeding the cells. Control and ZNF217 overexpressing cells (8 X 10^4^ cells) were seeded on each cover glass and incubated at 37 °C with 5% CO_2_ for 24 hours. Subsequently, the cells were washed twice with 1X PBS and fixed with 4% paraformaldehyde. Next, they were treated with Quenching solution (20 mM glycine, 20nM NH_4_Cl in PBS) for 10 mins and permeabilized with 0.1% Triton X in PBS for 10 mins at room temperature. For non-permeabilized IF staining, the treatment with 0.1% Triton-X/PBS was omitted. Cells were blocked with 4% Bovine Serum Albumin (BSA) in PBS for 1 hour and incubated with ERBB2 primary antibody (Proteintech #18299-I-AP, 1:300 in 4% BSA/PBS) overnight at 4 °C. Next day, the cells were washed gently with 1X PBS, blocked for 4% BSA/PBS, and incubated with FITC-anti rabbit secondary antibody (Proteintech #SA00003-2, 1:600 in 4% BSA/PBS) for 1 hour. Post-incubation, the cells were stained with Hoechst dye (Invitrogen #33342, 1:20000 in PBS) for 10 seconds, washed and mounted on glass slide using Prolong™ Gold Antifade mounting media. The images were taken at 63X using Zeiss LCM900 confocal microscope.

### RNA sequencing and data analysis

Flash frozen cell pellets were shipped on dry ice to Azenta Life Sciences. Three biological replicates were submitted for each sample. RNA extraction and RNA sequencing were performed at Azenta Life Sciences using their standardized methods. Proper quality check was included to ensure data quality. Preliminary data analysis was performed by Azenta Life Sciences as part of the package. Genes with an adjusted p-value < 0.05 and absolute log2 fold change >1 was considered significant. Volcano plot and heatmap were generated using the SRPlot platform^62^.

### Quantitative real-time PCR

Cells were harvested and RNA was isolated using RNA mini prep isolation kit (Zymo, #R1057). Extracted RNA was converted into cDNA using the iscript cDNA synthesis kit (Bio-Rad, #1708890). 20 ng of total cDNA was used for quantitative PCR (qPCR) for ERBB2 using Thermofisher SYBR Green master Mix kit. Cycling conditions was 95 °C for 2 minutes, 40 cycles of 95 °C for 15 seconds, 60 °C for 60 seconds, followed by melt curve analysis. The following ERBB2 primers were used to perform qPCR: forward primer: 5’-CATTTCTGCCGGAGAGCTT-3’ and reverse primer: 5’-GGTAACCTGTGATCTCTTCCAGA. Ct values of each sample were normalized to the Ct value for TATA-binding protein (TBP) and fold change was calculated using ΔΔCt method.

### In vivo mouse experiments

Animals were housed 2–5 per cage in temperature (22 ± 2 °C) and humidity (55% ± 15%) controlled rooms on a 12-hour light/dark cycle. Water and food were available to animals ad libitum. Animals are monitored daily for tumor growth and sacrificed when they meet euthanasia criteria outlined by NIH’s Guide for the Care and Use of Laboratory Animals. Luciferase-tagged OVCA420 and OVCA420-ZNF217 cells (10^7^ cells) were injected intraperitoneally into 6–8-week-old female Foxn1 nude mice. Mice injected with OVCA420-ZNF217 cells were randomized and assigned to three treatment groups: (1) vehicle (control); (2) 25 mg/kg TP-0903; and 2) 50 mg/kg TP-0903. An N=6 mice were used in each group. TP-0903 (Apex Bio #B5940) was freshly prepared each day by dissolving in corn oil at a concentration of 10 mg/ml and administered by oral gavage 5 days/week for 21 days. For IVIS imaging, mice were injected intraperitoneally with 150 mg/Kg D-luciferin in PBS, anesthetized, and bioluminescence signal (tumor burden) was acquired using the IVIS system. Signal intensity was quantified by Living Image analysis software (PerkinElmer). Tumor burden and survival was measured as outcomes. GraphPad prism was used to analyze the data and error bars shown are standard deviation. In vivo studies reported in this manuscript were conducted in accordance with a protocol that was approved by the Institutional Animal Care and Use Committee at the University of Maryland Baltimore County (protocol #1947). As ovarian cancer is a female-specific disease, we used only female age-matched mice were used.

### Quantification and data analysis

All In vitro assays were done in triplicates, 3 biological replicates with multiple technical replicates each time. All data points on graph represented as mean with standard deviation shown as error bars. Surface area covered by cells for in vitro assays was calculated using Image J/Fiji per field of image and normalized to the respective values in the control cells. For comparison between two groups, statistical significance was measured using Welch’s two-tailed *t* test. The IC50 values were calculated using GraphPad Prism by plotting drug concentration versus normalized cell viability response. Dose response curves were fitted using nonlinear regression with variable slope model and IC50 values were determined from the fitted curves with a 95% confidence interval. p≤0.05 (*), p≤0.01 (**) and p≤0.001 (***) were used to determine significance. An N =6 mice/group were used for in vivo experiments. Survival studies in mice were analyzed by Kaplan-Meier curves and log-rank (Mantel-Cox) tests. RNAseq was performed using biological triplicates. Distribution of read counts in libraries were examined before and after normalization. The original read counts were normalized to adjust for various factors such as variations of sequencing yield between samples. These normalized read counts were used to accurately determine differentially expressed genes. The overall similarity among samples were assessed by the euclidean distance between samples. This method was used to examine which samples are similar/different to each other and if they fit to the expectation from the experiment design. A comparison of gene expression between the groups of samples was performed using DESeq2. The Wald test was used to generate p-values and log2 fold changes. Genes with an adjusted p-value < 0.05 and absolute log2 fold change > 1 were called as differentially expressed genes. Significantly differentially expressed genes were clustered by their gene ontology and the enrichment of gene ontology terms was tested using Fisher exact test (GeneSCF v1.1-p2).

## DATA AVAILABILITY

All data supporting the findings of this study are available within the article and its supplementary file. RNAseq data for control and ZNF217 overexpressing OVCA420 and TYK-Nu cells are available in the Sequence Read Archive (SRA) database under accession code PRJNA1353133. RNAseq data from control and ZNF217 knockdown OVCA420 cells used in the manuscript are available in the Sequence Read Archive (SRA) database under accession code PRJNA1509651. There are no restrictions on data availability.

## ACKNOWLEDGEMENTS

A.P. has been supported by the Department of Defense (HT9425-23-1-0351 and HT9425-23-1-0232), National Institutes of Health (R03CA282712), funds from the Ovarian Cancer Alliance of Greater Cincinnati (PRIV0201 and PRIV0219), and UMGCCC American *Cancer* Society Institutional Research Grant – IRG-18-160-16. A.P was also supported by grants from the University of Maryland, Baltimore, Institute for Clinical & Translational Research (ICTR), which is funded in part by the National Center for Advancing Translational Sciences (NCATS) Clinical Translational Science Award (CTSA), UM1TR004926.

## AUTHOR CONTRIBUTIONS

Conceptualization — A.P.; Experimental design — M.J.P. and A.P.; Reagent development and data acquisition — M.J.P., J.H. and A.P.; Data analysis and interpretation — M.J.P. and A.P.; Writing original draft — M.J.P. and A.P.; Writing - review & editing — M.J.P., J.H. and A.P.; Funding acquisition and supervision — A.P.

## REFERENCES

1. Ortiz M, Wabel E, Mitchell K, et al. Mechanisms of chemotherapy resistance in ovarian cancer. Cancer Drug Resist 2022; 5: 304–316. 20220403. DOI: 10.20517/cdr.2021.147.

2. Wang L, Wang X, Zhu X, et al. Drug resistance in ovarian cancer: from mechanism to clinical trial. Mol Cancer 2024; 23: 66. 20240328. DOI: 10.1186/s12943-024-01967-3.

3. Havasi A, Cainap SS, Havasi AT, et al. Ovarian Cancer-Insights into Platinum Resistance and Overcoming It. Medicina (Kaunas) 2023; 59 20230310. DOI: 10.3390/medicina59030544.

4. Wardrup KC, Hoffman J, Pandya MJ, et al. ZNF217 promotes ovarian cancer progression by impacting multiple pivotal steps in the metastatic process. NPJ Precis Oncol 2025; 9: 392. 20251204. DOI: 10.1038/s41698-025-01153-8.

5. Stern DF. Tyrosine kinase signalling in breast cancer: ErbB family receptor tyrosine kinases. Breast Cancer Res 2000; 2: 176–183. 20000325. DOI: 10.1186/bcr51.

6. Kaneko S, Takasawa K, Asada K, et al. Mechanism of ERBB2 gene overexpression by the formation of super-enhancer with genomic structural abnormalities in lung adenocarcinoma without clinically actionable genetic alterations. Mol Cancer 2024; 23: 126. 20240611. DOI: 10.1186/s12943-024-02035-6.

7. Galogre M, Rodin D, Pyatnitskiy M, et al. A review of HER2 overexpression and somatic mutations in cancers. Crit Rev Oncol Hematol 2023; 186: 103997. 20230414. DOI: 10.1016/j.critrevonc.2023.103997.

8. Roskoski R, Jr. The ErbB/HER family of protein-tyrosine kinases and cancer. Pharmacol Res 2014; 79: 34–74. 20131120. DOI: 10.1016/j.phrs.2013.11.002.

9. Bai X, Sun P, Wang X, et al. Structure and dynamics of the EGFR/HER2 heterodimer. Cell Discov 2023; 9: 18. 20230213. DOI: 10.1038/s41421-023-00523-5.

10. Adam-Artigues A, Arenas EJ, Arribas J, et al. AXL - a new player in resistance to HER2 blockade. Cancer Treat Rev 2023; 121: 102639. 20231007. DOI: 10.1016/j.ctrv.2023.102639.

11. Adam-Artigues A, Arenas EJ, Martinez-Sabadell A, et al. Targeting HER2-AXL heterodimerization to overcome resistance to HER2 blockade in breast cancer. Sci Adv 2022; 8: eabk2746. 20220520. DOI: 10.1126/sciadv.abk2746.

12. Koziczak M and Hynes NE. Cooperation between fibroblast growth factor receptor-4 and ErbB2 in regulation of cyclin D1 translation. J Biol Chem 2004; 279: 50004–50011. 20040917. DOI: 10.1074/jbc.M404252200.

13. Penuel E, Akita RW and Sliwkowski MX. Identification of a region within the ErbB2/HER2 intracellular domain that is necessary for ligand-independent association. J Biol Chem 2002; 277: 28468–28473. 20020508. DOI: 10.1074/jbc.M202510200.

14. Nahta R. Molecular Mechanisms of Trastuzumab-Based Treatment in HER2-Overexpressing Breast Cancer. ISRN Oncol 2012; 2012: 428062. 20121122. DOI: 10.5402/2012/428062.

15. Carmona FJ, Montemurro F, Kannan S, et al. AKT signaling in ERBB2-amplified breast cancer. Pharmacol Ther 2016; 158: 63–70. 20151202. DOI: 10.1016/j.pharmthera.2015.11.013.

16. Cheng X. A Comprehensive Review of HER2 in Cancer Biology and Therapeutics. Genes (Basel) 2024; 15 20240711. DOI: 10.3390/genes15070903.

17. Hellstrom I, Goodman G, Pullman J, et al. Overexpression of HER-2 in ovarian carcinomas. Cancer Res 2001; 61: 2420–2423.

18. Caruso G, Weroha SJ and Cliby W. Ovarian Cancer: A Review. JAMA 2025; 334: 1278–1291. DOI: 10.1001/jama.2025.9495.

19. Zhang S, Pan Q and Wang K. Enhancing ovarian cancer treatment by synergistically targeting HER2 and PD-L1. Mol Ther Oncol 2025; 33: 200960. 20250311. DOI: 10.1016/j.omton.2025.200960.

20. Park J, Lee D, Kim S, et al. Prognostic implications of HER2 in ovarian cancer: Associations with homologous recombination deficiency and folate receptor alpha expression. Gynecol Oncol 2025; 203: 151–157. 20251104. DOI: 10.1016/j.ygyno.2025.10.029.

21. Bartha A and Gyorffy B. TNMplot: An enhanced platform for pharmacological target identification through cross-stage and pan-cancer gene expression analysis. Br J Pharmacol 2026; 183: 2648–2659. 20260320. DOI: 10.1111/bph.70390.

22. Gyorffy B. Discovery and ranking of the most robust prognostic biomarkers in serous ovarian cancer. Geroscience 2023; 45: 1889–1898. 20230301. DOI: 10.1007/s11357-023-00742-4.

23. Carney WP. HER2 status is an important biomarker in guiding personalized HER2 therapy. Per Med 2005; 2: 317–324. DOI: 10.2217/17410541.2.4.317.

24. De Cuyper A, Van Den Eynde M and Machiels JP. HER2 as a Predictive Biomarker and Treatment Target in Colorectal Cancer. Clin Colorectal Cancer 2020; 19: 65–72. 20200208. DOI: 10.1016/j.clcc.2020.02.007.

25. Loyez M, Lobry M, Hassan EM, et al. HER2 breast cancer biomarker detection using a sandwich optical fiber assay. Talanta 2021; 221: 121452. 20200730. DOI: 10.1016/j.talanta.2020.121452.

26. Harries M and Smith I. The development and clinical use of trastuzumab (Herceptin). Endocr Relat Cancer 2002; 9: 75–85. DOI: 10.1677/erc.0.0090075.

27. Hudis CA. Trastuzumab--mechanism of action and use in clinical practice. N Engl J Med 2007; 357: 39–51. DOI: 10.1056/NEJMra043186.

28. Maadi H, Soheilifar MH, Choi WS, et al. Trastuzumab Mechanism of Action; 20 Years of Research to Unravel a Dilemma. Cancers (Basel) 2021; 13 20210715. DOI: 10.3390/cancers13143540.

29. Tural D, Akar E, Mutlu H, et al. P95 HER2 fragments and breast cancer outcome. Expert Rev Anticancer Ther 2014; 14: 1089–1096. 20140626. DOI: 10.1586/14737140.2014.929946.

30. Scaltriti M, Rojo F, Ocana A, et al. Expression of p95HER2, a truncated form of the HER2 receptor, and response to anti-HER2 therapies in breast cancer. J Natl Cancer Inst 2007; 99: 628–638. DOI: 10.1093/jnci/djk134.

31. Jani JP, Finn RS, Campbell M, et al. Discovery and pharmacologic characterization of CP-724,714, a selective ErbB2 tyrosine kinase inhibitor. Cancer Res 2007; 67: 9887–9893. DOI: 10.1158/0008-5472.CAN-06-3559.

32. Vernieri C, Milano M, Brambilla M, et al. Resistance mechanisms to anti-HER2 therapies in HER2-positive breast cancer: Current knowledge, new research directions and therapeutic perspectives. Crit Rev Oncol Hematol 2019; 139: 53–66. 20190503. DOI: 10.1016/j.critrevonc.2019.05.001.

33. Zhang Q, Park E, Kani K, et al. Functional isolation of activated and unilaterally phosphorylated heterodimers of ERBB2 and ERBB3 as scaffolds in ligand-dependent signaling. Proc Natl Acad Sci U S A 2012; 109: 13237–13242. 20120625. DOI: 10.1073/pnas.1200105109.

34. Monje PV, Athauda G and Wood PM. Protein kinase A-mediated gating of neuregulin-dependent ErbB2-ErbB3 activation underlies the synergistic action of cAMP on Schwann cell proliferation. J Biol Chem 2008; 283: 34087–34100. 20080917. DOI: 10.1074/jbc.M802318200.

35. Stern DF. ERBB3/HER3 and ERBB2/HER2 duet in mammary development and breast cancer. J Mammary Gland Biol Neoplasia 2008; 13: 215–223. 20080503. DOI: 10.1007/s10911-008-9083-7.

36. Krig SR, Miller JK, Frietze S, et al. ZNF217, a candidate breast cancer oncogene amplified at 20q13, regulates expression of the ErbB3 receptor tyrosine kinase in breast cancer cells. Oncogene 2010; 29: 5500–5510. 20100726. DOI: 10.1038/onc.2010.289.

37. Oh DY and Bang YJ. HER2-targeted therapies - a role beyond breast cancer. Nat Rev Clin Oncol 2020; 17: 33–48. 20190923. DOI: 10.1038/s41571-019-0268-3.

38. Arteaga CL, Sliwkowski MX, Osborne CK, et al. Treatment of HER2-positive breast cancer: current status and future perspectives. Nat Rev Clin Oncol 2011; 9: 16–32. 20111129. DOI: 10.1038/nrclinonc.2011.177.

39. Okines A, Cunningham D and Chau I. Targeting the human EGFR family in esophagogastric cancer. Nat Rev Clin Oncol 2011; 8: 492–503. 20110405. DOI: 10.1038/nrclinonc.2011.45.

40. Bose R and Zhang X. The ErbB kinase domain: structural perspectives into kinase activation and inhibition. Exp Cell Res 2009; 315: 649–658. 20080815. DOI: 10.1016/j.yexcr.2008.07.031.

41. Nahta R. Pharmacological strategies to overcome HER2 cross-talk and Trastuzumab resistance. Curr Med Chem 2012; 19: 1065–1075. DOI: 10.2174/092986712799320691.

42. Hart JR and Vogt PK. Phosphorylation of AKT: a mutational analysis. Oncotarget 2011; 2: 467–476. DOI: 10.18632/oncotarget.293.

43. Littlepage LE, Adler AS, Kouros-Mehr H, et al. The transcription factor ZNF217 is a prognostic biomarker and therapeutic target during breast cancer progression. Cancer Discov 2012; 2: 638–651. 20120510. DOI: 10.1158/2159-8290.CD-12-0093.

44. Suarez CD, Wu J, Badve SS, et al. The AKT inhibitor triciribine in combination with paclitaxel has order-specific efficacy against Zfp217-induced breast cancer chemoresistance. Oncotarget 2017; 8: 108534–108547. 20170717. DOI: 10.18632/oncotarget.19308.

45. Yadav M, Sharma A, Patne K, et al. AXL signaling in cancer: from molecular insights to targeted therapies. Signal Transduct Target Ther 2025; 10: 37. 20250210. DOI: 10.1038/s41392-024-02121-7.

46. Corcoran RB, Ebi H, Turke AB, et al. EGFR-mediated re-activation of MAPK signaling contributes to insensitivity of BRAF mutant colorectal cancers to RAF inhibition with vemurafenib. Cancer Discov 2012; 2: 227–235. 20120116. DOI: 10.1158/2159-8290.CD-11-0341.

47. Li F, Huynh H, Li X, et al. FGFR-Mediated Reactivation of MAPK Signaling Attenuates Antitumor Effects of Imatinib in Gastrointestinal Stromal Tumors. Cancer Discov 2015; 5: 438–451. 20150211. DOI: 10.1158/2159-8290.CD-14-0763.

48. Tang Z, Li C, Kang B, et al. GEPIA: a web server for cancer and normal gene expression profiling and interactive analyses. Nucleic Acids Res 2017; 45: W98–W102. DOI: 10.1093/nar/gkx247.

49. Eleonora Teplinsky FM. EGFR and HER2: is there a role in ovarian cancer? Translational Cancer Research 2015; 4: 10. DOI: 10.3978/j.issn.2218-676X.2015.01.01.

50. Serra V, Scaltriti M, Prudkin L, et al. PI3K inhibition results in enhanced HER signaling and acquired ERK dependency in HER2-overexpressing breast cancer. Oncogene 2011; 30: 2547–2557. 20110131. DOI: 10.1038/onc.2010.626.

51. Antony J and Huang RY. AXL-Driven EMT State as a Targetable Conduit in Cancer. Cancer Res 2017; 77: 3725–3732. 20170630. DOI: 10.1158/0008-5472.CAN-17-0392.

52. Byers LA, Diao L, Wang J, et al. An epithelial-mesenchymal transition gene signature predicts resistance to EGFR and PI3K inhibitors and identifies Axl as a therapeutic target for overcoming EGFR inhibitor resistance. Clin Cancer Res 2013; 19: 279–290. 20121022. DOI: 10.1158/1078-0432.CCR-12-1558.

53. Taniguchi H, Yamada T, Wang R, et al. AXL confers intrinsic resistance to osimertinib and advances the emergence of tolerant cells. Nat Commun 2019; 10: 259. 20190116. DOI: 10.1038/s41467-018-08074-0.

54. Leconet W, Chentouf M, du Manoir S, et al. Therapeutic Activity of Anti-AXL Antibody against Triple-Negative Breast Cancer Patient-Derived Xenografts and Metastasis. Clin Cancer Res 2017; 23: 2806–2816. 20161206. DOI: 10.1158/1078-0432.CCR-16-1316.

55. Goyette MA, Cusseddu R, Elkholi I, et al. AXL knockdown gene signature reveals a drug repurposing opportunity for a class of antipsychotics to reduce growth and metastasis of triple-negative breast cancer. Oncotarget 2019; 10: 2055–2067. 20190312. DOI: 10.18632/oncotarget.26725.

56. Wei J, Sun H, Zhang A, et al. A novel AXL chimeric antigen receptor endows T cells with anti-tumor effects against triple negative breast cancers. Cell Immunol 2018; 331: 49–58. 20180514. DOI: 10.1016/j.cellimm.2018.05.004.

57. Schoumacher M and Burbridge M. Key Roles of AXL and MER Receptor Tyrosine Kinases in Resistance to Multiple Anticancer Therapies. Curr Oncol Rep 2017; 19: 19. DOI: 10.1007/s11912-017-0579-4.

58. Liu L, Greger J, Shi H, et al. Novel mechanism of lapatinib resistance in HER2-positive breast tumor cells: activation of AXL. Cancer Res 2009; 69: 6871–6878. 20090811. DOI: 10.1158/0008-5472.CAN-08-4490.

59. Halmos B and Haura EB. New twists in the AXL(e) of tumor progression. Sci Signal 2016; 9: fs14. 20161004. DOI: 10.1126/scisignal.aai7619.

60. Caro AA, Gordun Peiro A, Hadadi E, et al. A fast-progressing orthotopic ovarian cancer model reveals synergistic antitumor effects of AXL-targeting nanobodies and Olaparib. Gynecol Oncol Rep 2025; 62: 101998. 20251125. DOI: 10.1016/j.gore.2025.101998.

61. Wiederschain D, Wee S, Chen L, et al. Single-vector inducible lentiviral RNAi system for oncology target validation. Cell Cycle 2009; 8: 498–504. 20090225. DOI: 10.4161/cc.8.3.7701.

62. Tang D, Chen M, Huang X, et al. SRplot: A free online platform for data visualization and graphing. PLoS One 2023; 18: e0294236. 20231109. DOI: 10.1371/journal.pone.0294236.

