## Supplemental Data for "ZNF217 promotes receptor tyrosine kinase plasticity and AXL-ERK dependency in ovarian cancer"

**SUPPLEMENTAL TABLES AND TABLE LEGENDS**

| Target | Source | Catalog number | Dilution |
|---|---|---|---|
| ERBB2 | Proteintech | 18299-I-AP | 1:1000 (WB)<br>1:300 (IF) |
| ZNF217 | Invitrogen | 720352 | 1:1000 |
| ERBB3 | Cell signaling | 12708S | 1:1000 (WB) |
| EGFR | Proteintech | 18986-I-AP | 1:1000 (WB) |
| pERB2(Tyr877) | Proteintech | 82762-11-RR | 1:1000 (WB) |
| p-AKT(S473) | Proteintech | 66444-1-IG | 1:1000 (WB) |
| p-AKT(T308) | Proteintech | 81232-10-RR | 1:1000 (WB) |
| PDK1 | Proteintech | 18262-1-AP | 1:1000 (WB) |
| AKT | Proteintech | 10176-2-AP | 1:1000 (WB) |
| PTEN | Proteintech | 22034-I-AP | 1:1000 (WB) |
| FGFR1 | Proteintech | 60325-1-IG | 1:1000 (WB) |
| FGFR4 | Proteintech | 81069-1-RR | 1:1000 (WB) |
| AXL | Proteintech | 84659-1-RR | 1:1000 (WB) |
| p-AXL(Tyr779) | Proteintech | 82816-4-RR | 1:1000 (WB) |
| ERK 1/2 | Proteintech | 11257-I-AP | 1:1000 (WB) |
| pERK1/2 | Proteintech | 80031-1-RR | 1:1000 (WB) |
| Beta-Actin | Sigma | A228 | 1:10000 (WB) |
| FITC-anti rabbit | Proteintech | SA00003-2 | 1:600 (IF) |

**Table S1**. List of antibodies, their source and dilutions used in this study.

### SUPPLEMENTAL FIGURES AND FIGURE LEGENDS

**Fig. S1**

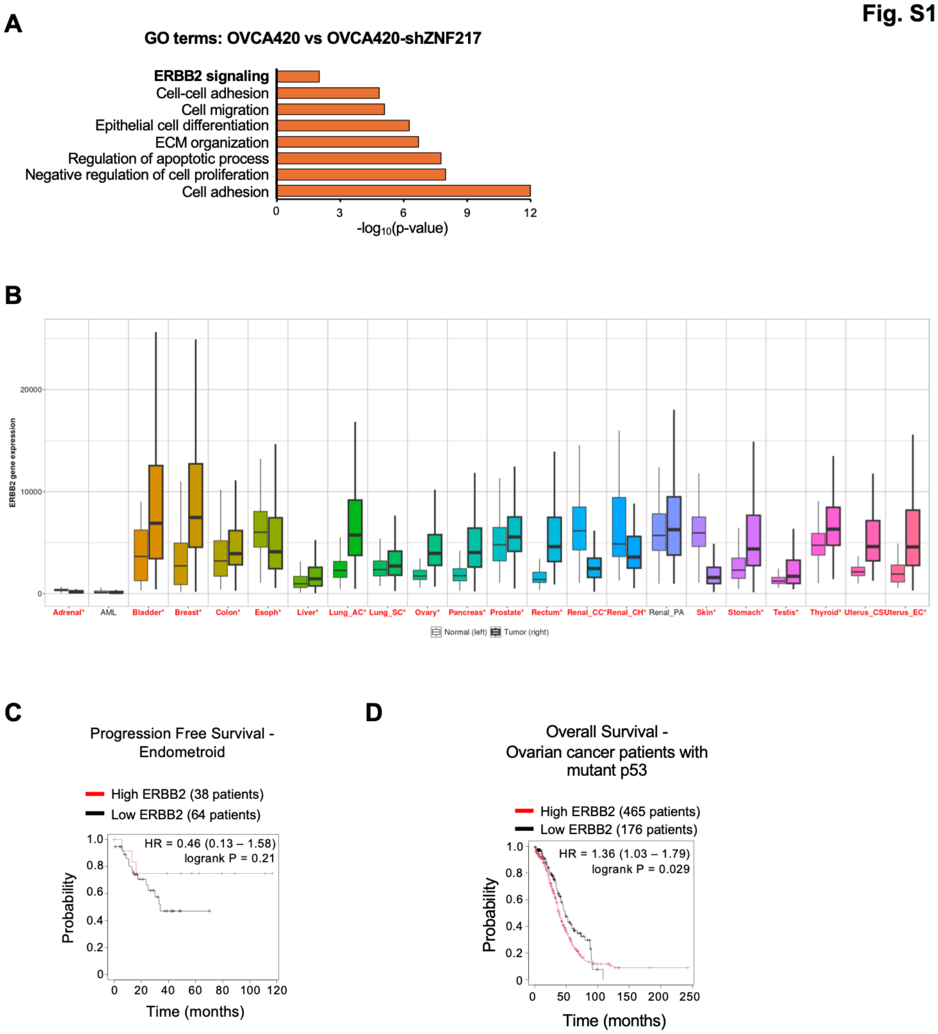


**Figure S1. ERBB2 is overexpressed in ovarian cancer and associated with poor prognosis.** **(A)** Gene ontology terms that are altered upon ZNF217 knockdown in OVCA420 cells. **(B)** ERBB2 expression is altered in many cancers including ovarian cancer. Data retrieved from TNMplot. **(C)** ERBB2 overexpression does not impact the progression free survival (PFS) in endometroid ovarian tumors. **(D)** ERBB2 overexpression causes a reduction in overall survival (OS) in mutant p53 driven ovarian cancer.

Fig. S2

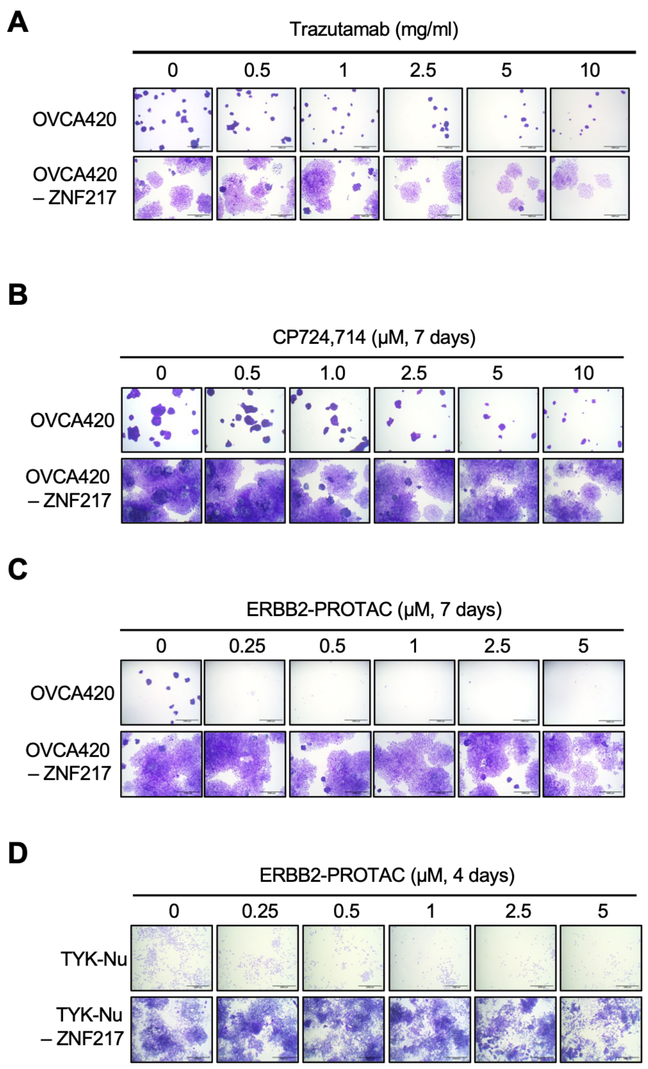


**Figure S2. ZNF217-high ovarian cancer cells are more resistant to ERBB2-targeting therapeutics (A)** Representative crystal violet staining showing OVCA420-ZNF217 cells are more resistant to trastuzumab (7 days treatment). **(B)** Representative crystal violet staining showing OVCA420-ZNF217 cells are more resistant to CP724,714 (7 days treatment). **(C)** Representative crystal violet staining showing OVCA420-ZNF217 cells are more resistant than control OVCA420 cells to ERBB2-PROTAC (7 days treatment). **(D)** Representative crystal violet staining showing TYK-Nu-ZNF217 cells are more resistant than control TYK-Nu cells to ERBB2-PROTAC (4 days treatment).

Fig. S3

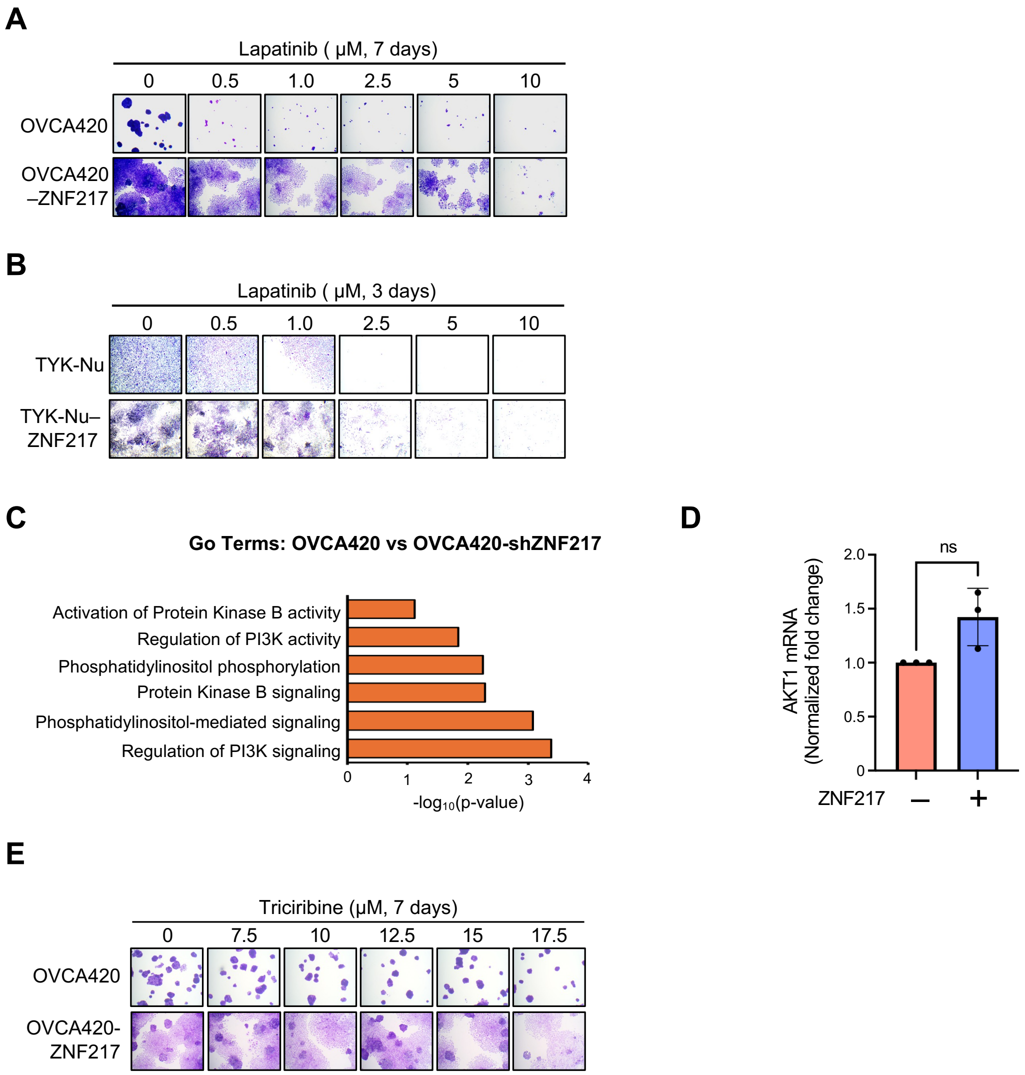


**Figure S3. ZNF217-mediated resistance to anti-ERBB2 therapeutics is not dependent on EGFR and AKT signaling pathway. (A)** Representative crystal violet staining showing OVCA420-ZNF217 cells are more resistant to lapatinib (7 days treatment). **(B)** Representative crystal violet staining showing TYK-Nu-ZNF217 cells are more resistant to lapatinib (4 days treatment). **(C)** Protein Kinase B signaling related Gene Ontology terms are significantly enriched in the differentially expressed gene sets in ZNF217 knockdown OVCA420 cells. **(D)** RT-qPCR analysis showing no change in AKT mRNA levels upon ZNF217 overexpression in OVCA420 cells. **(E)** Representative crystal violet staining showing that sensitivity of OVCA420 cells to long-term triciribine treatment (7 days) is not impacted by ZNF217 overexpression.

Fig. S4

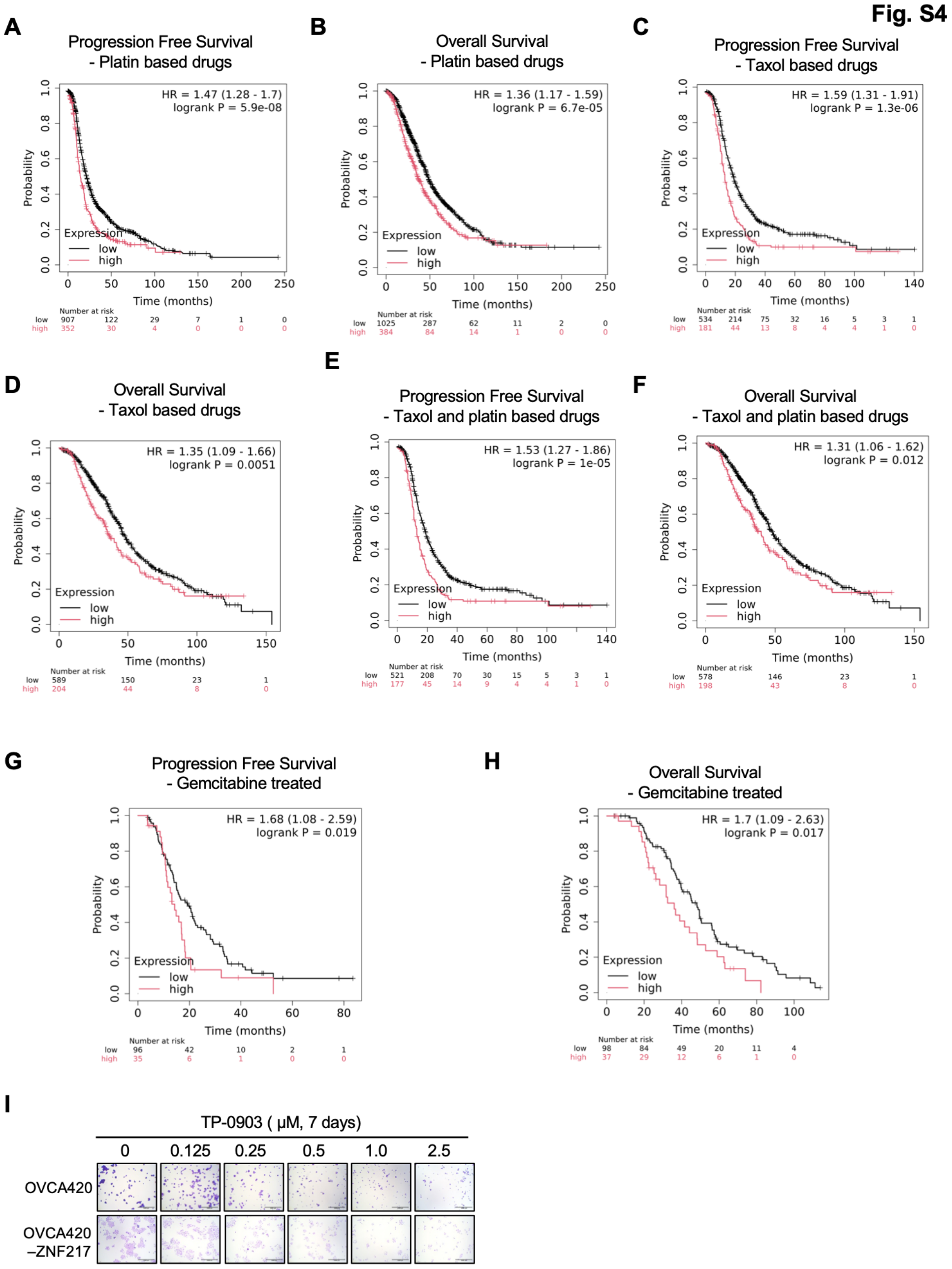


**Figure S4. AXL overexpression in ovarian tumors is associated with increased resistance to multiple chemotherapeutic agents.** (**A** and **B**) Clinical data from Kmplotter database show that patients with AXL-high ovarian tumors exhibit lower **(A)** progression free survival and **(B)** overall survival when treated with platin-based drugs. (**C** and **D**) Clinical data from Kmplotter database show that patients with AXL-high ovarian tumors exhibit lower **(C)** progression free survival and **(D)** overall survival when treated with taxol-based drugs. (**E** and **F**) Clinical data from

Kmplotter database show that patients with AXL-high ovarian tumors exhibit lower **(E)** progression free survival and **(F)** overall survival when treated with a combination of platin- and taxol-based drugs. **(G** and **H**) Clinical data from Kmplotter database show that patients with AXL-high ovarian tumors exhibit lower **(G)** progression free survival and **(H)** overall survival when treated with gemcitabine. **(I)** Representative crystal violet staining showing OVCA420-ZNF217 cells are more sensitive to the AXL inhibitor, TP-0903 (7 days treatment).

Fig. S5

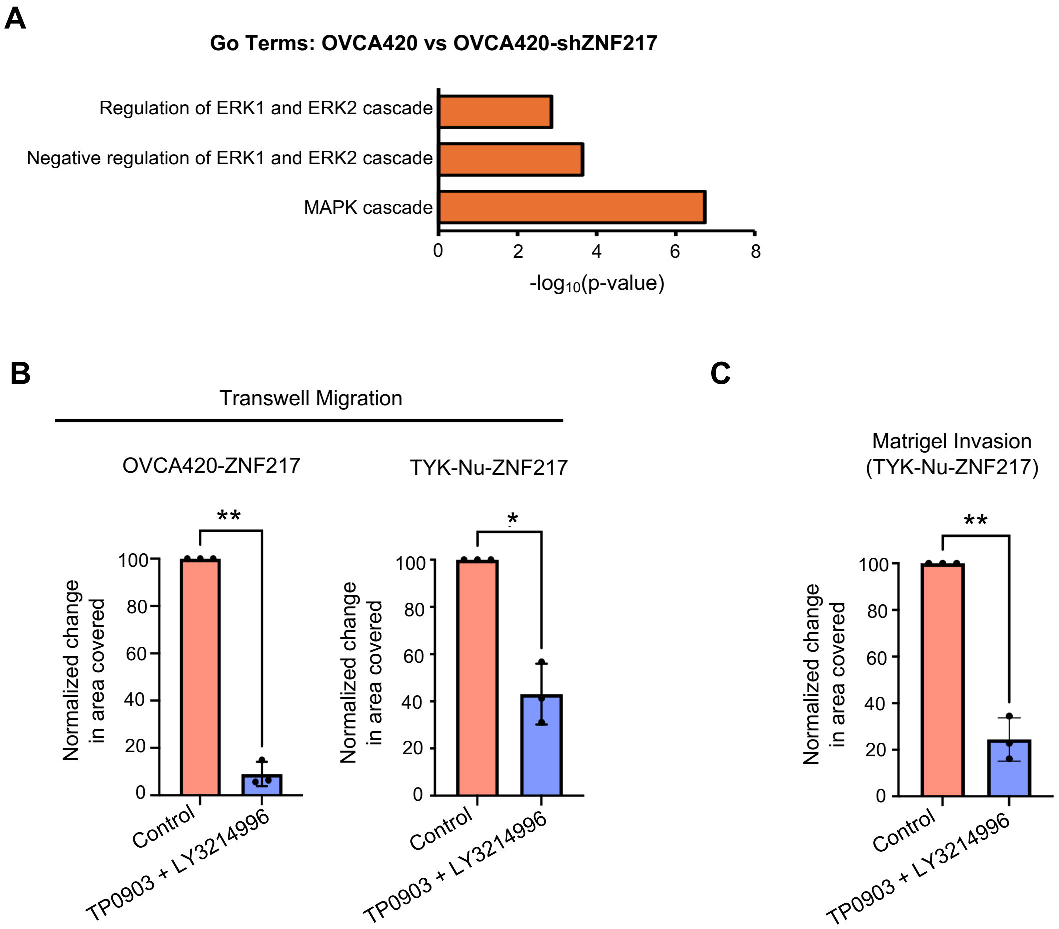


**Figure S5. MAPK signaling is critical for ZNF217 driven metastatic phenotypes in ovarian cancer cells. (A)** MAPK signaling related Gene Ontology terms are significantly enriched in the differentially expressed gene sets in ZNF217 depleted OVCA420 cells. **(B)** Effect of combined inhibition of AXL using TP-0903 (100 nM) and ERK using LY3214996 (1 μM) on cell migration in OVCA420-ZNF217 and TYK-Nu-ZNF217 cells using transwell migration assay. **(C)** Effect of combined inhibition of AXL using TP-0903 (100 nM) and ERK using LY3214996 (1 μM) on the ability if TYK-Nu-ZNF217 cells to invade through the matrigel.
